# Target gene regulatory network of miR-497 in angiosarcoma

**DOI:** 10.1101/2023.09.24.559218

**Authors:** Annaleigh Benton, Emma Terwilliger, Noah M. Moriarty, Bozhi Liu, Ant Murphy, Hannah Maluvac, Mae Shu, Lauren E. Gartenhaus, Nimod D. Janson, Claire M. Pfeffer, Sagar M. Utturkar, Elizabeth I. Parkinson, Nadia A. Lanman, Jason A. Hanna

## Abstract

Angiosarcoma (AS) is a vascular sarcoma that is highly aggressive and metastatic. Due to its rarity, treatment options for patients are limited, therefore more research is needed to identify possible therapeutic vulnerabilities. We previously found that conditional deletion of *Dicer1* drives AS development in mice. Given the role of DICER1 in canonical microRNA (miRNA) biogenesis, this suggests that miRNA loss is important in AS development. After testing miRNAs previously suggested to have a tumor-suppressive role in AS, microRNA-497-5p (miR-497) suppressed cell viability most significantly. We also found that miR-497 overexpression led to significantly reduced cell migration and tumor formation. To understand the mechanism of miR-497 in tumor suppression, we identified clinically relevant target genes using a combination of RNA-sequencing data in an AS cell line, expression data from AS patients, and target prediction algorithms. We validated miR-497 direct regulation of CCND2, CDK6, and VAT1. One of these genes, VAT1, is an understudied protein that has been suggested to promote cell migration and metastasis in other cancers. Indeed, we find that pharmacologic inhibition of VAT1 with the natural product Neocarzilin A reduces AS migration. This work provides insight into the mechanisms of miR-497 and its target genes in AS pathogenesis.

## INTRODUCTION

Angiosarcoma (AS) is an aggressive, highly metastatic endothelial cell cancer. The five-year survival rate of AS is only 30%, and patients with metastatic disease have a particularly poor prognosis, with a median survival time of only 12 months^1,2^. Because AS is a rare cancer, much remains unknown about the disease. Commonly mutated genes have been identified, including *KDR*, *TP53*, POT1, *PTPRB,* and *PIK3CA*^3–6^. Recent studies have investigated novel potential treatments, including combined VEGFR and MAPK inhibition, PI3K inhibitors, propranolol, and TKIs combined with immunotherapy^3,7–9^. However, elucidating crucial genetic drivers and other therapeutic targets will be imperative for additional progress. Approximately 30% of AS patients present with *de novo* metastatic disease, and half of those diagnosed with localized AS will develop metastatic disease later^2, 10,11^. This high incidence of metastasis highlights the need for the development of therapeutics that can prevent the occurrence of metastatic dissemination.

MicroRNAs (miRNAs) are small RNAs that regulate gene expression through binding to target gene transcripts, resulting in mRNA degradation or translational repression^12^. MiRNAs undergo several essential processing steps mediated by biogenesis machinery, including DICER1, which cleaves precursor miRNAs to form mature miRNAs. Canonically, this step is required for the formation of mature, functional miRNAs, and disruption of this pathway can result in global mature miRNA loss and lead to disease^13^.

Most biological processes are influenced by miRNA activity, therefore their dysregulation can contribute to various pathologies, such as cancer^14–18^. In cancer, miRNAs can function as both oncogenes and tumor suppressors^19,20^, which is largely dependent on the genes they regulate and the cellular context. Given the roles of miRNAs in cancer and their ability to regulate a multitude of transcripts, miRNAs themselves have been implicated as a potential therapeutic targets, through either inhibition of oncogenic miRNAs or replacement of tumor-suppressive miRNAs^21,22^.

Although the importance of miRNAs in cancer is well-defined, and widespread dysregulation of miRNAs have been reported in soft tissue sarcomas^23^, there is much to discover regarding the roles of miRNAs in AS. Previously it has been suggested that miR-210^24^, miR-214^25^, miR-340^26^, and miR-497^27^ may elicit tumor-suppressing phenotypes in AS. Furthermore, *DICER1* mutations have been reported in patient AS samples, suggesting that global miRNA perturbations may be an important component of AS pathogenesis^28^. In addition, we previously demonstrated biallelic loss of *Dicer1* in endothelial cells drives AS tumor formation in mice^29,30^. Given the essential role DICER1 serves in miRNA biogenesis, these findings suggest that miRNAs loss plays a critical role in the development of AS, therefore warranting more investigation into their role in disease progression. In the present study, we aim to further characterize the role of miRNAs in AS. These findings will further define the mechanism of tumor suppressive miRNAs in AS through the regulation of critical target genes.

## RESULTS

### MiR-497 significantly reduces cell viability in AS cell line panel

We have previously found that the loss of miRNA processing machinery, *Dicer1*^29^, in endothelial cells leads to angiosarcoma development in mice, suggesting that miRNA loss may be an important factor in AS pathogenesis. However, much remains unknown about the contribution of individual miRNAs in AS. We previously found that miR-23 target genes were enriched in *Dicer1* knockout mouse angiosarcomas^29^. Additionally, some miRNAs have been implicated to have tumor-suppressive properties in AS, such as miR-210^24^, miR-214^25^, miR-340^26^, and miR-497^27^. We sought to select a miRNA of interest to further study from these five candidates. To evaluate the effect of overexpression of each miRNA, miRNA mimics were transfected into a panel of mouse (Fig. 1A) and human (Fig. 1B) cell lines, including MS1, HMEC-1 (endothelial), EOMA (hemangioendothelioma), SVR, ADC106, and AS5 (AS). Successful mimic transfection was validated by qRT-PCR for each mimic compared to the negative control (NC) mimic (Fig. 1C, Supplementary Fig. S1A-D). Among the candidate miRNA mimics tested, miR-497 was the only candidate that significantly suppressed cell viability in all the tested AS cell lines (Fig. 1A-C). Furthermore, miR-497 mimic transfection increased apoptosis in AS and hemangioendothelioma cell lines (Fig. 1D). Given the observed phenotypes across this cell line panel, we further characterized the tumor suppressive role of miR-497 in AS.

**Figure 1:**
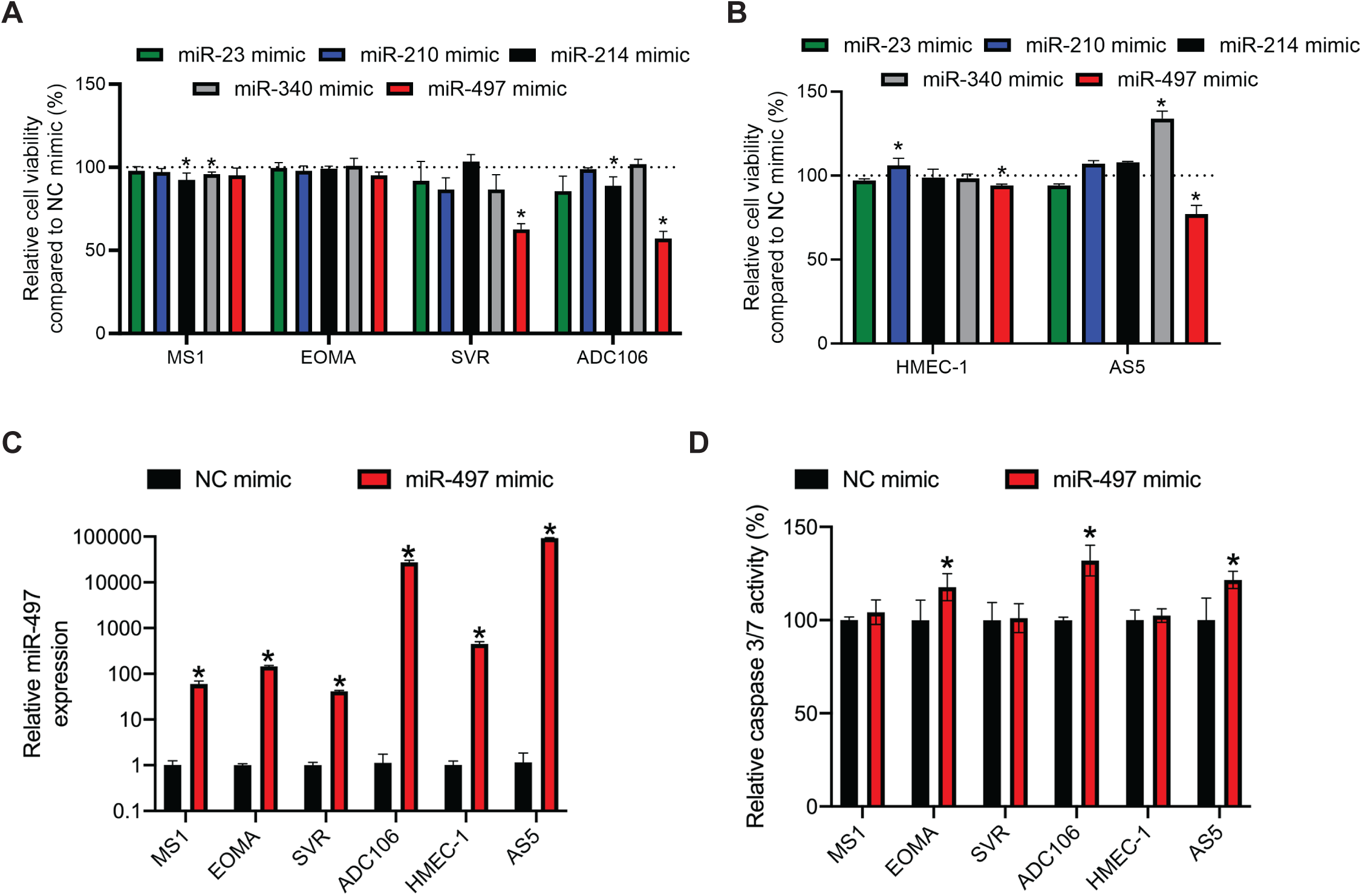
Transfection of miR-497 mimic inhibits cell viability and induces apoptosis in angiosarcoma cell lines. **(A)** Normalized cell viability in mouse and **(B)** human cell lines 72 hours post-transfection with indicated miRNA mimics relative to negative control (NC) mimic. **(C)** Relative expression of miR-497-5p by qRT-PCR in cells as in (A). **(D)** Relative caspase 3/7 activity in cell line panel 72 hours after NC or miR-497 mimic transfection in indicated cell line, * p<0.05.

### Constitutive expression of miR-497 inhibits cell viability, migration, and tumor **formation**

To interrogate the effect of miR-497 overexpression in AS, the pre-miR-497 was cloned into a lentiviral vector and constitutively expressed in ADC106 cells (Fig. 2A). Expression of mature miR-497 was significantly increased compared to empty control cells (Fig. 2B) and Clover was similarly expressed (Fig. 2C). Similar to the mimic transfections, constitutive expression of miR-497 reduced cell viability (Supplementary Fig. 2A) and population doubling times. Compared to empty control (Fig. 2D), suggesting that miR-497 overexpression impacts tumor cell proliferation. In agreement with previous studies in ISO-HAS cells^27^, transwell migration assays revealed that cell migration was also significantly reduced in pre-miR-497 cells (Fig. 2E).

**Figure 2:**
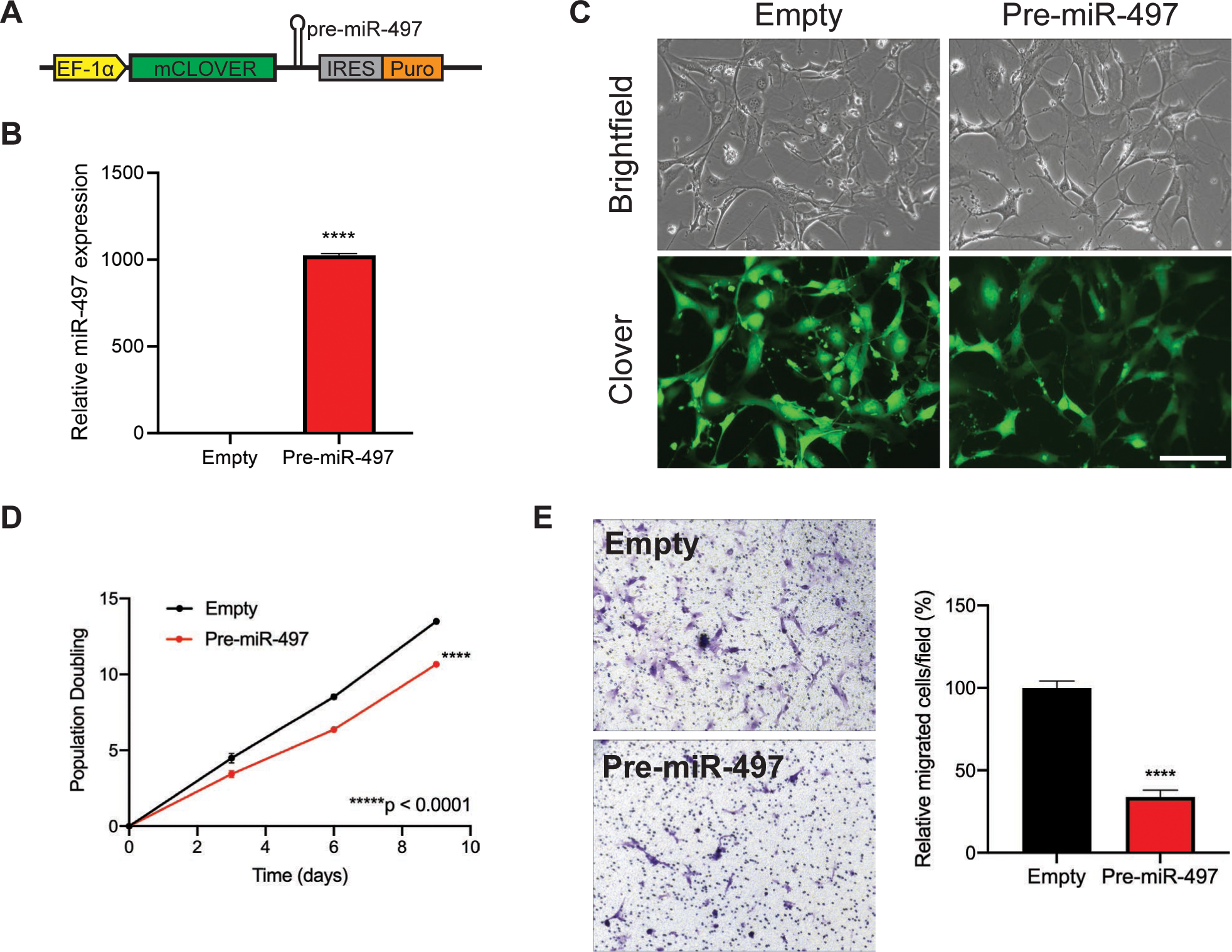
Lentiviral miR-497 expression reduces proliferation and migration in AS cells. **(A)** Schematic of pSIN-Clover-Puro lentiviral vector used to overexpress pre-miR-497. **(B)** Relative expression of miR-497 by qRT-PCR in ADC106 cells stably transduced with pSIN-Clover-Empty or pre-miR-497. **(C)** Brightfield and Clover images of ADC106 cells stably transduced with pSIN-Clover-Empty control or pre-miR-497, scale bar 150 μm. **(D)** Population doubling assay and **(E)** representative images of transwell migration assays with quantified relative migrated cells of ADC106 empty and pre-miR-497 cells. ****p<0.0001.

To determine the role of miR-497 in tumor formation, pre-miR-497 and control ADC106 cells were subcutaneously injected into the flank of immune-compromised (NRG) mice. Expression of pre-miR-497 significantly reduced tumor formation in mice compared to control such that palpable tumors were not detected in pre-miR-497 injected mice (Fig. 3A-B). Upon dissection, small masses were observed in the pre-miR-497 cells indicating miR-497 expression significantly reduced tumor volume (Fig. 3C) and final tumor mass (Fig. 3D). Congruent with reduced proliferation in miR-497 expressing cells, the percent of Ki-67 positive cells was significantly decreased in pre-miR-497 tumors compared to empty control tumors (Fig. 3E-F). To assess the role miR-497 plays in patient survival, using publicly available sarcoma patient data from the TCGA, we found that high miR-497 expression correlates with improved patient survival (Supplementary Fig. 2B)^31^. Taken together, this data suggests that miR-497 can attenuate several cancer phenotypes, including proliferation, cell migration, and tumor formation.

**Figure 3:**
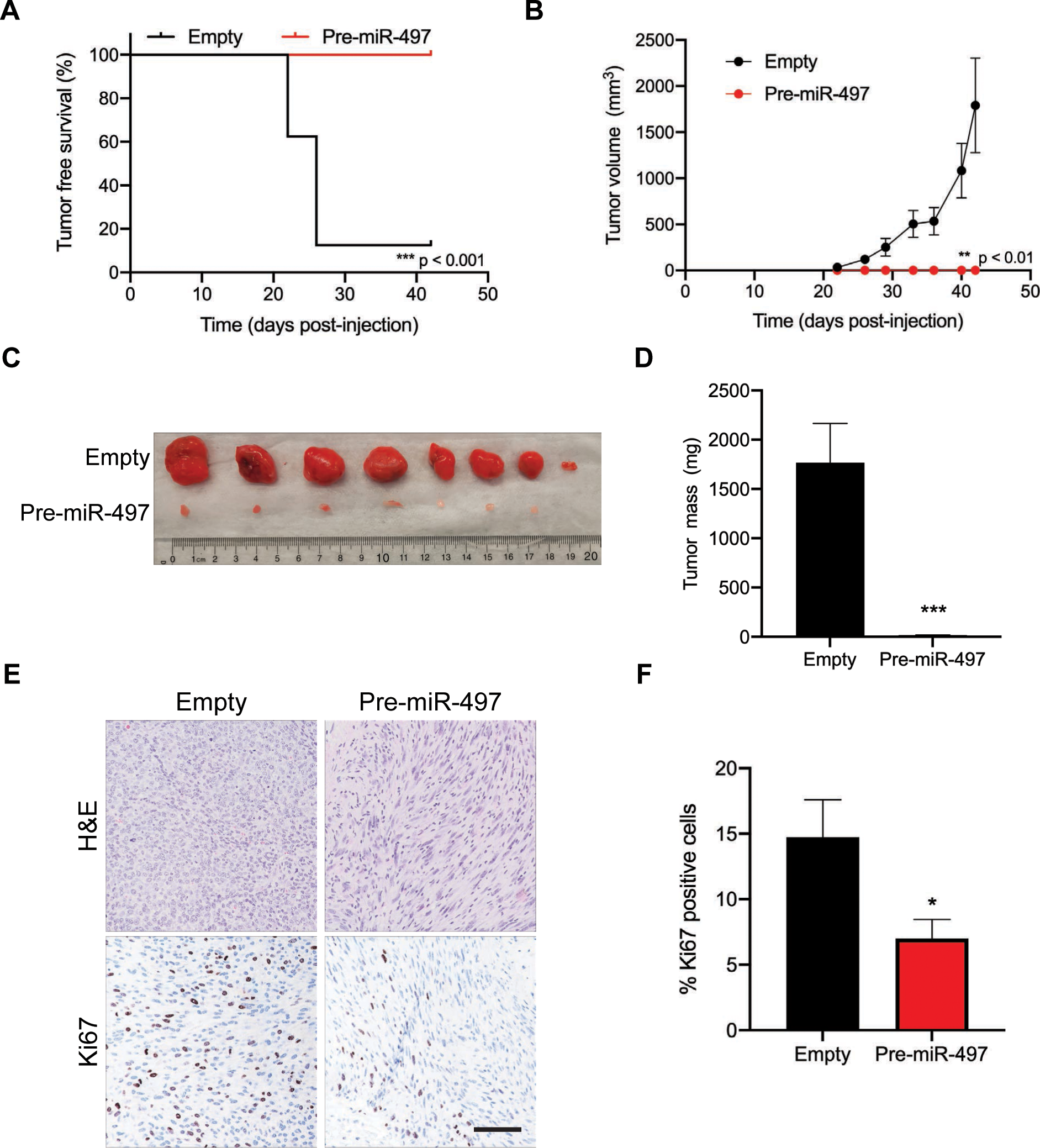
MiR-497 overexpression ablates AS tumor formation. **(A)** Tumor-free survival in mice subcutaneously injected with pSIN-Clover-Empty control and pre-miR-497 ADC106 cells (n=8). **(B)** Subcutaneous tumor growth of pSIN-Clover-Empty and pre-miR-497 ADC106 cells, represented as mean ± S.E.M. **(C)** Final tumor images and **(D)** tumor mass of pSIN-Clover-Empty and pre-miR-497 ADC106 cells. **(E)** Representative images of H&E and Ki67 stained tumors, scale bar is 100 μm. **(F)** Quantification of Ki67 positive cells in allografts. *p<0.05, **p<0.01, ***p<0.001.

### Candidate target genes of miR-497

Phenotypes observed as a result of miRNA overexpression are largely achieved through the regulation of target gene transcripts. Therefore, to elucidate the mechanism of miR-497 tumor suppression, we sought to identify the miR-497 target gene candidates and the enriched pathways in cells overexpressing miR-497. We transfected ADC106 cells with miR-497 or NC mimics and conducted RNA sequencing (Fig. 4A and Supplementary Fig. 3A). Importantly, predicted miR-497 target genes were significantly downregulated based on GSEA, ENRICHR analysis, and CDF plots (Fig. 4B&C, Supplementary Fig. 3B, Supplementary Table S1).

**Figure 4:**
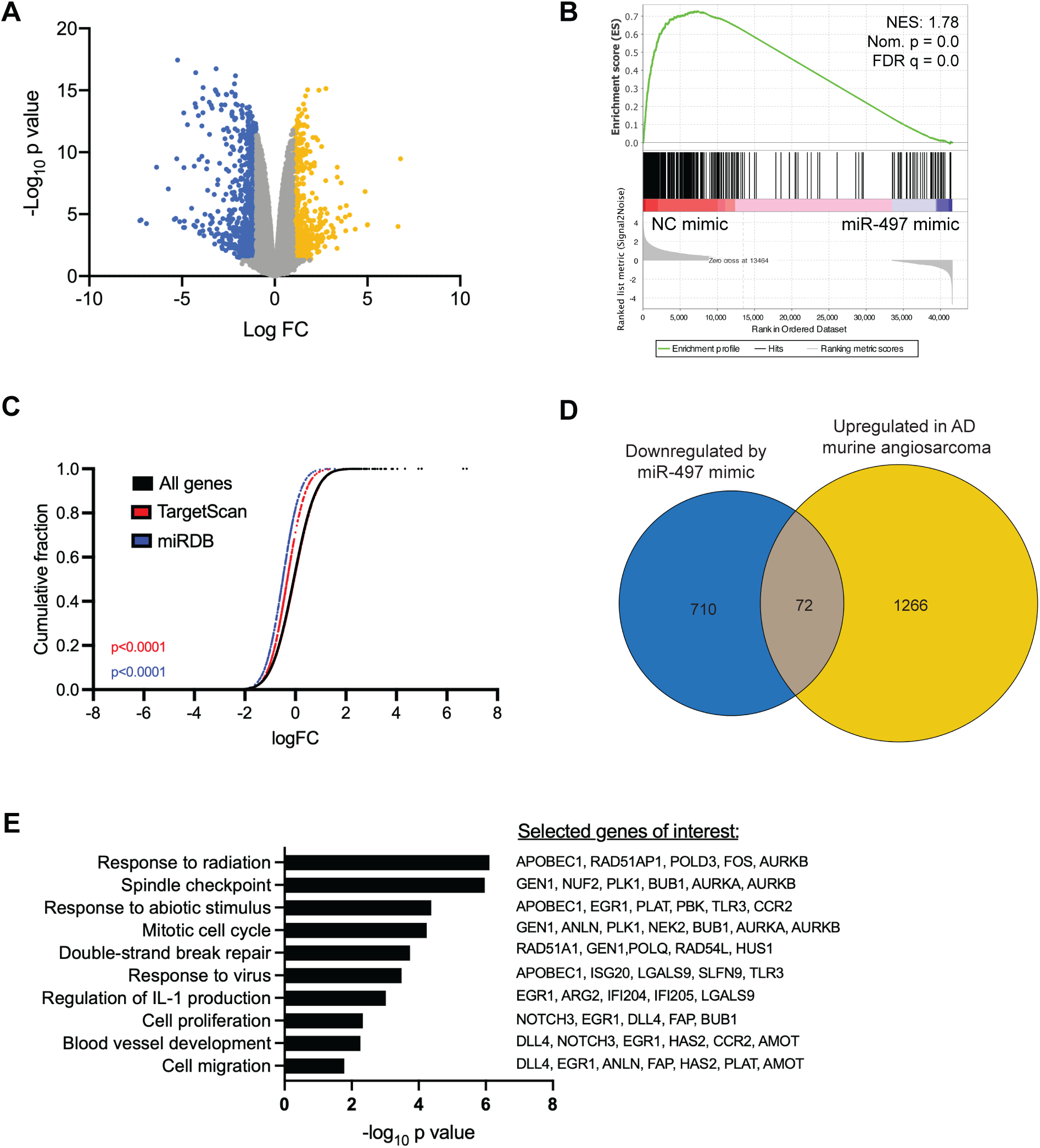
Target gene regulatory network of miR-497. **(A)** Volcano plot of the Log_10_ of the p value versus the log fold change (FC) in mRNA expression in miR-497 mimic transfected versus NC mimic transfected ADC106 cells. Genes with p < 0.05 and log FC > 1 (yellow) or log FC < −1 (blue). **(B)** Gene Set Enrichment Analysis (GSEA) demonstrates enrichment of miR-497 predicted target genes with normalized enrichment score (NES) of 1.78, nominal p value of 0.0, and FDR q value of 0.0. **(C)** Alterations of gene expression changes plotted as a cumulative distribution frequency (CDF). Significant downregulation of predicted murine miR-497 target genes based on Targetscan (1,169 genes, red) and miRDB (375 genes, blue) compared to all genes upon miR-497 mimic transfection. Similarity of miR-497 predicted target distributions to the total distribution was tested with the one-sided Kolmogorov–Smirnov test. **(D)** Euler diagram of number of genes increased in *aP2-Cre;Dicer^cKO^*(AD) tumors vs aorta with FDR < 0.05 and log FC > 1 (1,338 genes) and are down with miR-497 FDR < 0.05 and log FC < −1 (782 genes). **(E)** Gene ontology of the common 72 genes from (D) with selected genes of interest from each enriched term.

Gene ontology (GO) terms related to regulation of cell adhesion, extracellular matrix organization, negative regulation of migration, and angiogenesis were upregulated in miR-497 transfected cells likely due to secondary effects of miR-497 (Supplementary Fig. 3C and Supplementary Table S2). Whereas response to virus, regulation of phosphorylation, and apoptotic process GO terms were enriched among the downregulated genes (Supplementary Fig. 3D and Supplementary Table S3). To determine the miR-497 target genes potentially involved in AS, GO was performed on the 72 overlapping genes downregulated in the miR-497 transfected cells and upregulated in *aP2-Cre;Dicer1^cKO^* (AD^cKO^)^30^ murine AS tumors (Fig. 4D). Terms such as spindle checkpoint (PLK1, AURKB, AURKA), mitotic cell cycle (GEN1, PLK1, NEK2), and blood vessel development (DLL4, NOTCH3, EGR1) were amongst the top enriched pathways (Fig. 4E and Supplementary Table S4).

### Validation of miR-497 target gene candidates

To identify direct target genes of miR-497 most clinically relevant to AS, we compared the genes downregulated by miR-497 in ADC106 cells to those reported to be upregulated in patient angiosarcomas^32^, and predicted targets of miR-497 from TargetScan^33^. Based on these criteria, we identified four genes, *Cdk6, Ccnd2, Dll4*, and *Vat1* for further study (Fig. 5A). In addition, 14 genes were significantly downregulated by miR-497 overexpression and were overexpressed in patient tumors but were not predicted targets of miR-497. These genes may highlight indirect consequences of miR-497 overexpression or genes with non-canonical regulation. Some genes in this group promote cell proliferation, tumorigenesis and/or metastasis, such as FOS, SFRP2, MAPK3, and TNFAIP6.

**Figure 5:**
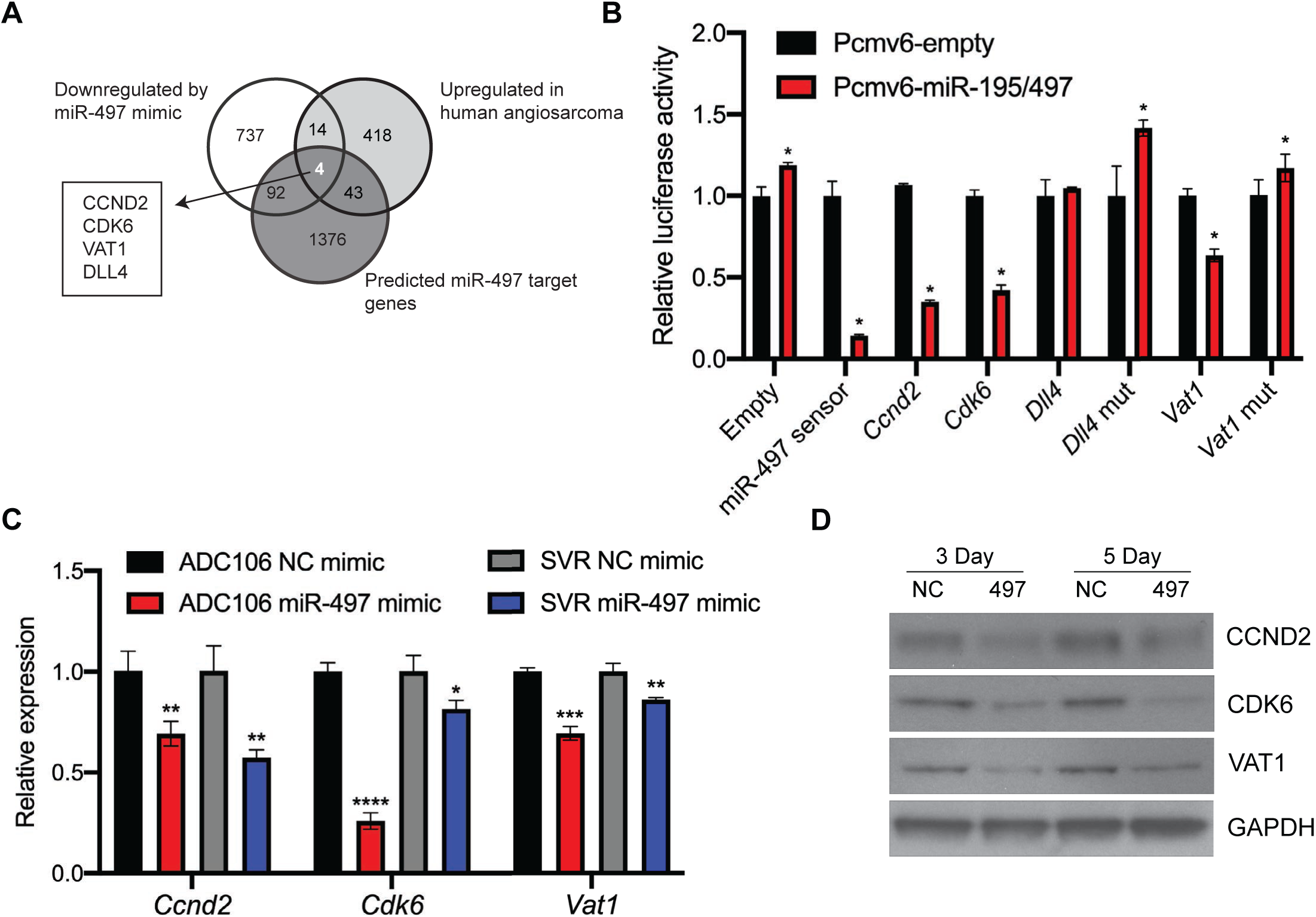
Validation of miR-497 target genes. **(A)** Venn diagram of the genes downregulated by miR-497 mimic with p<0.05, and log FC<-1 (847 genes), upregulated in human AS tumors with p< 0.05 and log FC>3.5 (479 genes) and have conserved, predicted miR-497 sites annotated in TargetScan (1,515 genes). **(B)** Luciferase activity in 293T cells co-transfected with pre-miR-195/497 or control vector, and empty psiCHECK2 reporter, miR-497 sensor positive control, mouse wild-type 3’ UTRs of indicated genes, or 3’ UTRs with mutated miR-497 sites. Renilla/Firefly luciferase ratio normalized to empty reporter (no miR-497). **(C)** Relative expression of miR-497 target genes by qRT-PCR analysis 3 days after NC or miR-497 mimic transfection in indicated cells, *p<0.05, **p<0.01, ***p<0.001, ****p<0.0001. **(D)** Immunoblot analysis of ADC106 cells post transfect with NC or miR-497 mimic at indicated time points.

To test specificity of the miR-497 regulation of *Ccnd2*, *Cdk6*, *Dll4,* and *Vat1*, 3’ UTR dual luciferase assays were conducted to validate these candidates as functional target genes of miR-497. The 3’ UTRs were cloned into the psicheck2 dual luciferase reporter plasmid. This confirmed that *Ccnd2*, *Cdk6*, and *Vat1* 3’ UTRs are indeed regulated by miR-497 (Fig. 5B). Mutation of the first three nucleotides of the miR-497 binding sites in the 3’ UTR of *Vat1* abolished the ability of miR-497 to regulate it, therefore confirming the specificity of miR-497 regulation of the *Vat1* transcript. miR-497 direct regulation of the human 3’ UTR of *CDK6* was also confirmed, and mutagenesis of the miR-497 binding site abolished the observed regulation (Supplementary Fig. 4A-D). Interestingly, regulation of the 3’ UTR of *Dll4* was not observed (Fig. 5A). The specificity of miR-497 regulation of the *Ccnd2* transcript has been previously demonstrated^34^. Based on this data, we confirm that miR-497 does indeed functionally regulate target gene candidates, *Vat1*, *Ccnd2*, and *Cdk6*. We were unable to confirm that miR-497 regulates the 3’ UTR of *Dll4*, therefore we did not include it in further studies. To further validate *Vat1*, *Ccnd2*, and *Cdk6* as target gene transcripts of miR-497, we performed qRT-PCR and Western blots on NC and miR-497 mimic transfected cells. We observed that miR-497 mimic transfection decreased the expression of these target gene transcripts in ADC106 and SVR cells (Fig. 5C). Additionally, miR-497 expression reduced protein levels for each at 3- and 5-days post transfection in the ADC106 cells (Fig. 5D). Taken together, this data validates that miR-497 directly regulates *Vat1*, *Ccnd2*, and *Cdk6* in AS cells.

### *Vat1* mediates cell migration in AS cell lines

We chose to further characterize the role of VAT1 in AS, as VAT1 is relatively understudied. VAT1 has been previously implicated in synaptic vesicle transport, mitochondrial fusion, oxidoreductase function, and cancer cell migration^35–42^. Given that miR-497 reduced cell migration in AS cells and can regulate VAT1, we hypothesized that VAT1 may promote cell migration in AS.

We generated ADC106 and SVR cells with doxycycline-inducible shRNAs directed against *VAT1* or *Renilla* control (Ren). Both *VAT1* shRNAs A and B reduced *Vat1* expression by qRT-PCR and immunoblot (Supplementary Fig. 5A-C). In both cell lines, a more robust reduction of VAT1 protein was observed with shA (Supplementary Fig. 5B-C). Furthermore, we found that the shA modestly reduced cell migration by transwell migration assay, whereas knockdown by shB did not significantly reduce migration (Supplementary Fig. 5D-E). This difference in migration effects may be due to the residual VAT1 protein and differences in knockdown efficiencies between shA and shB.

Due to the observed knockdown efficiency issues and residual VAT1 protein, we also utilized a pharmacologic approach to inhibit VAT1. Neocarzilin A (NCA) is a natural product that was previously shown to inhibit VAT1 via chemical proteomic studies, and was also shown to significantly reduce cell migration in cancer cell lines^35^. Thus, we synthesized NCA as previously described^35^ (see Supplementary Methods). To test NCA efficacy, we utilized the human breast cancer cell line, MDA-MB-231 as these cells have previously been found to be sensitive to NCA^35^. We initially determined the IC_50_ of NCA in the MDA-MB-231, MS1, SVR, and ADC106 cells (Fig. 6A). The IC_50_ values were between 1-2 µM which is consistent with previous studies^35^. Of note, the normal endothelial, MS1, cells had the highest IC_50_. Next, we found that NCA inhibited cell migration in MDA-MB-231 breast cancer cells as previously shown^35^. Furthermore, NCA significantly decreased cell migration in the ADC106 and SVR AS cell lines (Fig. 6B-C). Thus, while the shRNA knockdown of *Vat1* had modest effects on migration, the inhibition of VAT1 with NCA significantly reduced AS cell migration. Additional studies on the specificity of NCA for VAT1, and the *in vivo* efficacy of NCA and VAT1 inhibition will be explored in future studies.

**Figure 6:**
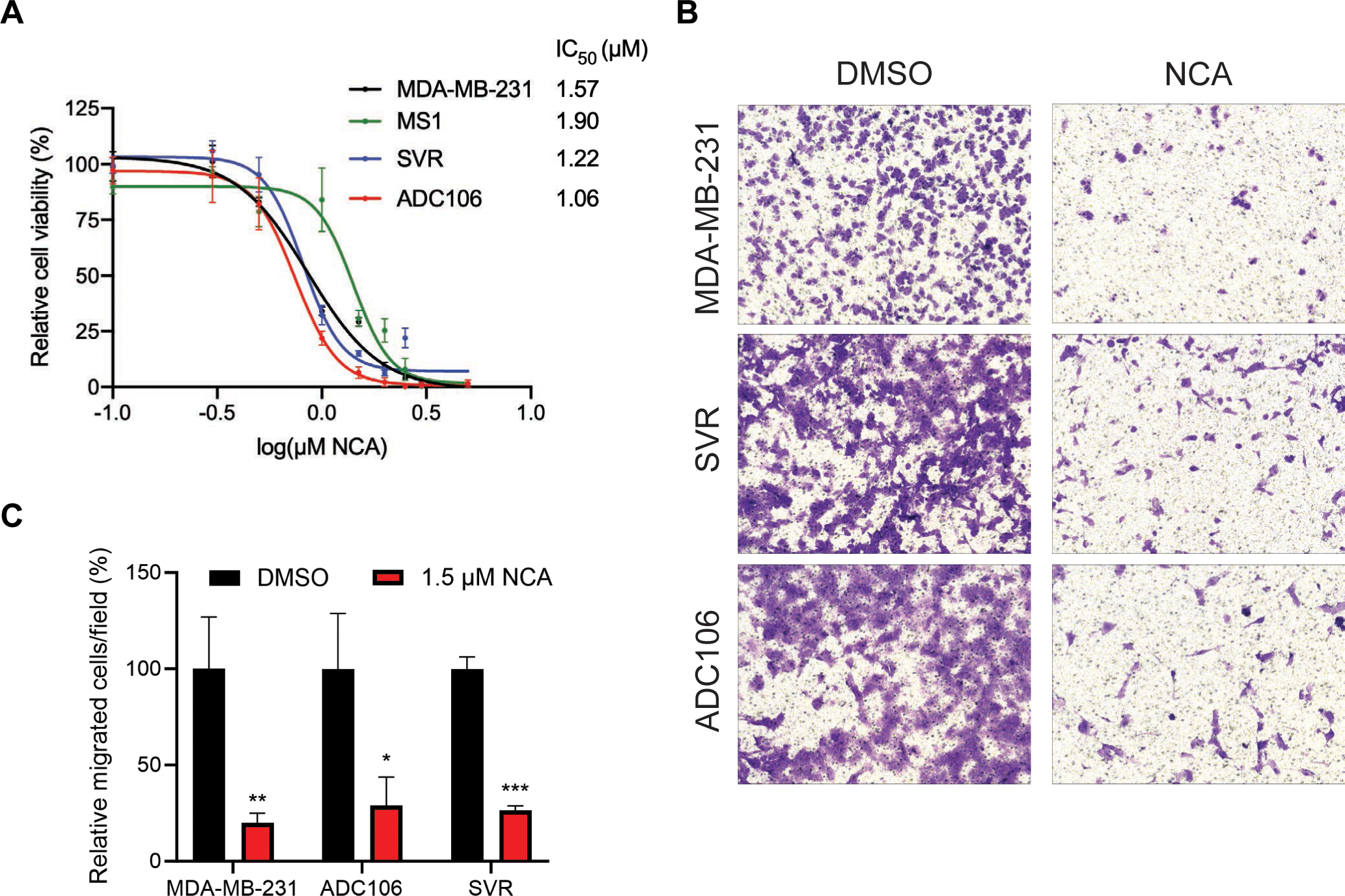
Pharmacologic inhibition of miR-497 target gene, VAT1, by natural product, NCA, inhibits cell migration. **(A)** Cell viability curve based on Cell Titer Glo analysis of MS1, SVR, ADC106, and MDA-MB-231 cells, 72 hours after treatment with indicated concentrations of VAT1 inhibitor, Neocarzilin A (NCA). **(B)** Transwell migration assays of SVR, ADC106, and MDA-MB-231 cells pretreated with 1.5 µM NCA for 24 hours. **(C)** Quantification of the number of migration cells as from (B). *p<0.05, **p<0.01, ***p<0.001.

## DISCUSSION

Angiosarcoma (AS) is a rare, and aggressive endothelial cancer that is not well understood. Resources to study AS are limited, therefore developing models to study the disease is crucial for making novel discoveries. We have previously developed mouse models to recapitulate AS, including models driven by oncogenic KRAS and conditional *Dicer1* deletion in endothelial cells^29,30,43^. Because *Dicer1* is essential for microRNA (miRNA) biogenesis, this suggests that loss of mature miRNAs is responsible for this tumor development, however, other non-canonical *Dicer1* could be attributed as well. Nonetheless, these findings provide evidence that the regulation of miRNAs is an important component of AS pathogenesis.

MiRNAs are short non-coding RNAs that bind to the 3’ UTR of target gene transcripts based on sequence complementarity, resulting in transcript degradation or translational repression^12^. A miRNA seed sequence can be complementary to sites present in many mRNAs; therefore, a single miRNA may be predicted to regulate thousands of target gene transcripts. Because miRNAs are master regulators of many genes, they can possess the ability to target multiple oncogenic drivers or tumor-suppressors at once. Thus, individual miRNAs can be oncogenic or tumor-suppressive depending on their repertoire of target gene transcripts in a particular cellular context^19^. Because of their potential to modulate multiple oncogenic pathways, miRNAs have been pursued as promising therapeutics for cancer, through miRNA replacement or inhibition^44,45^. As miRNA therapeutics are being developed, one of the challenges will be identifying relevant miRNA candidates and miRNA targets for specific cancer types^22^. Therefore, understanding the roles of individual miRNAs in AS will be critical to pursue the possibility of miRNA therapeutics in the future.

The roles of some miRNAs have already been described in AS, including miR-210, miR-214, miR-340, and miR-497, each of them being characterized as tumor suppressors^24–27^. To select a single candidate to study further, we tested how overexpression of these miRNAs affected cell viability in a cell line panel. Interestingly, only miR-497 suppressed cell viability across all three AS cell lines we tested, and some candidates did not significantly suppress cell viability in any AS cell lines tested. The molecular heterogeneity of AS may explain this phenomenon^46^. However, because miR-497 most consistently reduced cell viability in several AS cell lines, this suggests that miR-497 serves a common role across a broad range of AS models, despite the heterogeneity of molecular drivers in AS. Therefore, we characterized miR-497 role in AS and determined target genes that may contribute to this phenotype.

MiR-497 is a member of the miR15/16/195/424/497 gene family, sharing a common seed sequence (AGCAGCA), and therefore predicted to regulate the same genes. Like its other family members, miR-497 is generally described as a tumor-suppressive miRNA in a variety of cancers and has been shown to be significantly downregulated in patient tumors^47–49^, including AS^27^. MiR-497 has been previously studied in AS, and inhibits cell viability, invasion, and tumor formation partially through regulation of potassium channel, KCa3.1 (*KCCN4*)^27^. Through transient and constitutive overexpression of miR-497, we have further defined the tumor suppressive roles of miR-497 in other AS cell lines, confirming that miR-497 reduced cell viability, proliferation, and migration. Our results also found that miR-497 overexpression was highly effective in reducing tumor formation, highlighting the potential therapeutic efficacy of miR-497. To further elucidate the mechanism of miR-497 we characterized the regulatory network of miR-497 target genes in AS.

MiR-497 was previously reported to elicit tumor-suppressive phenotypes through the regulation of the target gene transcript, *KCCN4* (KCa3.1) in AS^27^. We did not observe a significant reduction of *Kccn4* in our RNA-sequencing data. Thus, KCCN4 may be a context specific target gene of miR-497. Nonetheless, miR-497 overexpression led to tumor-suppressing phenotypes similarly in AS cell lines we tested. To further characterize clinically relevant target genes of miR-497, we identified genes significantly reduced by transient miR-497 overexpression, and cross referenced these genes with those significantly overexpressed in a patient data set and predicted target genes of miR-497 from TargetScan^33^. From this analysis, we identified four candidate target genes of miR-497 that are overexpressed in patient tumors, *Vat1* (Vesicle amine transport protein 1), *Ccnd2* (Cyclin-D2), *Cdk6* (Cyclin-dependent kinase 6), and *Dll4* (Delta like canonical notch ligand 4), and were able to validate three of the four original candidates, *Vat1*, *Ccnd2*, and *Cdk6*. Downregulation of cell cycle genes, *Ccnd2* and *Cdk6*, may explain the reduction in cell viability and proliferation observed from miR-497 overexpression. Additionally, miR-497 overexpression appears to significantly influence cell cycle differences in cells, given that the gene ontology (GO) term “regulation of cell cycle” was enriched in our RNA-sequencing data with miR-497 overexpression. We were able to validate *Ccnd2* and *Cdk6* as target genes of miR-497. Additionally, *Cdk6* has also been validated as a target gene of miR-497 in other cancers^50^. Dll4 is a ligand in the notch pathway, and regulation of this gene may play a role in angiogenesis-related signaling^51,52^. Additionally, GO terms “angiogenesis” and “blood vessel development” are enriched in the miR-497 RNA-sequencing data. However, we were unable to experimentally confirm that *Dll4* is directly regulated by miR-497.

*Vat1* has not yet been validated as a miR-497 target gene, nor has its role been studied in AS. VAT1 has been previously implicated in synaptic vesicle transport, mitochondrial fusion, oxidoreductase function, and cancer cell migration^35,37,39–42^. Previous reports have reported a pro-migratory role of VAT1 in glioblastoma, breast cancer, and hepatocellular carcinoma^35–37^. Our findings indicate that VAT1 knockdown had a modest effect on cell migration. Additionally, we also inhibited VAT1 with the natural product neocarzilin A (NCA) from *Streptomyces carzinostaticus*^53^. NCA is the only reported inhibitor of VAT1 to date and has been shown to reduce cell viability and migration in cancer cell lines^35^. Because VAT1 is overexpressed in AS, we hypothesized that NCA may be efficacious for reducing cell viability and cell migration. We found that NCA treatment significantly suppressed cell migration in AS cell lines, alluding to its potential as an anti-metastatic drug. NCA was much more efficacious in impairing cell migration compared to knockdown of *Vat1*. Since NCA is proposed to be a covalent inhibitor of VAT1^35^, it may produce greater affects through altered interactions with VAT1 binding proteins than simply reducing protein levels with shRNA-knockdown. However, since the mechanism through which VAT1 inhibition by NCA produces an antimigratory effect is unknown, further research is required to answer these questions. However, this provides additional evidence to support further characterization of VAT1 functions in cell migration and in vivo testing of NCA efficacy in cancer models.

In this study, we further characterized the role of miR-497 in AS, showing that it reduces cell viability, migration, and tumor formation. Because of these observed phenotypes, miR-497 may be an attractive candidate for miRNA therapy in AS. We found that miR-497 may elicit these phenotypes through regulation of several direct target genes overexpressed in AS, including *Ccnd2*, *Cdk6*, and *Vat1*. Finally, inhibition of VAT1 through the natural product, NCA, significantly impairs cell migration, which may indicate that that NCA could be a potential therapy to impair metastasis.

## MATERIALS AND METHODS

### Cell culture

Cells were obtained from the following sources: HMEC-1 (CRL-3243, American Type Culture Collection [ATCC], Manassas, VA, USA), MS1 (ATCC, CRL-2279), SVR (ATCC, CRL-2280), EOMA (M. Roussel, St. Jude Children’s Research Hospital), ADC106 (derived from our lab as previously described^30^), HEK-293T (M. Kazemian, Purdue University), MDA-MB-231 (M. Wendt, Purdue University), AS5 (E. Dickerson, University of Minnesota^54,55^). HMEC-1 cells were maintained in MCDB131 (15100CV Corning, NY, USA) supplemented with 10 ng/ml EGF (23626, R&D Systems, Minneapolis, MN, USA), 1 μg/ml hydrocortisone (0219456901, MP Biomedicals, Irvine, CA, USA), 1× Glutamax (35050, Gibco/Thermo Fisher Scientific, Waltham, MA, USA), and 10% FBS (SH3091003, Hyclone, Logan, UT, USA). AS5 cells were maintained in EBM-2 Basal Medium supplemented with EGM-2 SingleQuots (CC-3162, Lonza, Walkersville, MD, USA). The EOMA, MS1, SVR, MDA-MB-231, HEK-293T and ADC106 cells were maintained in DMEM (SH30243, Hyclone) with 10% FBS (SH30910.03, Hyclone), 1× antibiotic-antimycotic penicillin, streptomycin, and amphotericin B (PSA) (A5955, Sigma-Aldrich, St. Louis, MO, USA) and incubated at 37 °C in 5% CO2. STR analysis for the AS5, ADC106, MS1, and SVR cells performed by IDEXX (Supplementary Table S5). Cell viability was determined using the Cell Titer Glo Assay (G7570, Promega, Madison, WI, USA) and apoptosis was determined using the Caspase 3/7 Glo Assay (G8090, Promega) measuring luminescence with a BioTek Synergy 2 (BioTek/Agilent, Winooski, VT, USA). Cells were treated with NCA following its synthesis detailed in supplementary methods. Transwell assays were performed with transwell permeable supports (3422, Corning), between 2.5 x 10 and 1.0 x 10^5^ cells suspended in serum free DMEM, and DMEM supplemented with 10% FBS was used as a chemoattractant. Non-migrated cells were removed from the inner chamber with a cotton swab, migrated cells were crystal violet stained, imaged, and quantified. At least three independent fields per sample were used for quantification of images. Cells were transfected with 12.5 nM miRCURY LNA miRNA mimics: Negative Control (NC), miR-23a, miR-210, miR-214, miR-340 or miR-497 (339306, Qiagen) following the manufacturer’s instruction with Lipofectamine RNAiMax (13778075; Invitrogen/Thermo Fisher Scientific, Waltham, MA, USA).

### RNA and gene expression

Total RNA was prepared using the *Quick*-RNA Miniprep Kit (R1054, Zymo, Irvine, CA, USA) according to the manufacturer’s instructions. cDNA was generated using High-Capacity RNA to cDNA kit (4387406, Applied Biosystems, Waltham, Massachusetts, USA) and the miRCURY cDNA kit (339340, Qiagen, Germantown, MD, USA) for miRNA expression analysis. qRT-PCR was performed with SYBR and LNA miRCURY primers detailed in Supplementary Table S6. Relative expression by qRT-PCR was quantified using the delta-delta CT method normalized to *Gapdh*. Mature miRNA expression was normalized to snRNA U6. RNA-seq was performed on total RNA from ADC106 cells transfected with NC or miR-497 mimic. mRNA library prep (poly A enrichment) and RNA-seq performed by Novogene on an Illumina NovaSeq platform. Each sample contained at least 30 million paired end reads with length of 150 bp. Raw reads were trimmed using fastp^56^ (version 0.19.5). Quality trimmed reads were aligned to the mouse (GRCm38) genome using the STAR^57^ aligner (version 2.5.4b). Reads aligned to each gene feature were quantified using the featureCounts^58^ program from the Subread package (version 1.6.1). A differential expression (DE) analysis was performed using edgeR^59^ (version 3.32.1) and genes with p value ≤ 0.05 and log_2_(fold-change) > 1 or < −1 were denoted as significant. Differentially expressed genes were analyzed for gene ontology enrichment using the Database for Annotation, Visualization and Integrated Discovery (DAVID)^60^. Functional classification of terms or pathways were analyzed based on biological process gene ontology terms (BP_FAT) with the exclusion of redundant terms.

### Molecular cloning, viral transduction, and luciferase assays

pSIN-Clover-Puro and pSIN-Clover-miR-497-Puro were generated from the modified pSIN-Puro backbone utilizing pre-miR-497 cloned from the pMSCV-PIG-miR-497/miR-195 (Addgene #64236, a gift from Joshua Mendel^17^). Doxycycline-inducible shRNAs targeting Renilla and Vat1 were generated in LT3GEPIR^61^ (gift from Johannes Zuber, Addgene plasmid # 111177). The 97-mer shRNA sequences were ordered from IDT, PCR amplified, and assembled into the LT3GEPIR plasmid as previously described^61^. Further molecular cloning details and plasmid generation information provided in Supplementary Information. Lentivirus was produced in HEK-293T cells, by transfecting lentiviral vectors (10 µg), packaging (7.5 µg of psPAX2 Addgene #12260, a gift from Didier Trono), and envelope (3 µg of pMD2.G, Addgene #12259, a gift from Didier Trono) vectors using FuGENE6 (11 815 091 001, Roche, Basel, Switzerland). Virus was harvested from cell media 2- and 3-days post-transfection, filtered through 0.45 µm syringe filters, and applied to cells. 48 hours post transduction cells were selected and maintained in 2 µg/mL puromycin. Luciferase assays were performed by co-transfecting HEK-293T cells with 100 ng pCMV6-empty or pCMV6-miR-497/195 with 10 ng of psiCHECK2 reporter plasmids using FuGENE6 (E2691; Promega) and performing the Dual-Luciferase Reporter Assay System (E1910; Promega) after 72 h as previously described^18^.

### Subcutaneous tumor allografts

Immune-compromised (NOD.Cg-Rag1^tm1Mom^ Il2rg^tm1Wjl/SzJ^ (NRG) #007799, The Jackson Laboratory) mice were subcutaneously injected with 5.0 x 10^5^ ADC106 pSIN-Clover-Empty and pSIN-Clover-pre-miR-497 cells into the flank. Tumor volume was recorded twice weekly, and mice were euthanized when a tumor volume of 2 cm^3^ was reached. All mouse studies were reviewed and approved by the Institutional Animal Care and Use Committee at Purdue University.

### Immunohistochemistry

H&E staining performed by the Purdue Histology Research Laboratory following standard methods^62^. For Ki67 staining, antigen retrieval was performed by placing slides in 1x Tris EDTA buffer at 95 °C for 20 minutes. The slides were stained at room temp with primary antibody (Ki67 #275R-15, Cell Marque, Rocklin, California, USA) diluted 1:1000 for 30 min, then conjugated with the secondary (Goat anti-Rabbit ImmPRESS HRP, #MP-7451 Vector Labs, Newark, CA, USA) for an additional 30 min. DAB (ImmPACT DAB Substrate HRP#SK-4105, Vector Labs) was applied for 5 minutes and rinsed with water.

### Immunoblots

Protein lysates of cells and tissues were prepared in RIPA lysate buffer as previously described^18^. Protein concentrations were determined by the BCA Protein Assay Kit (23225, Pierce, Rockford, IL, USA). Equally loaded lysates were resolved by SDS-PAGE using NuPAGE 4-12% Bis-Tris Gel (NP0321BOX, Invitrogen/Thermo Fisher Scientific, Waltham, MA, USA) and transferred to Immobilon-P Transfer Membrane (IPVH00005, Merck Millipore, Burlington, MA). Blots were probed overnight at 4 °C with primary antibodies detailed in Supplementary Table S8. Following washing, the membranes were then probed with the species-appropriate secondary antibody. After washing the membranes again, protein expression was visualized using chemiluminescence Luminol reagent (SC-2048, Santa Cruz Biotechnology, Dallas, TX, USA) and film exposure (45-001-508 Amersham Hyperfilm ECL Film, Cytiva, Marlborough, MA, USA) or Azure c600 imager (Azure Biosystems, Dublin, CA, USA).

### Statistics

Statistical analyses were performed using Prism Version 9 (Graph Pad Software, Inc., San Diego, CA, USA). All results are expressed as the mean ± SD unless stated otherwise. Pairwise comparisons were performed with a two-tailed, unpaired Student’s t-test. All p values < 0.05 were considered significant.

## Acknowledgements

This work was supported in part by the Purdue Institute for Cancer Research (P30CA023168), a Purdue University Ross Fellowship (AB), the Purdue University Department of Biological Sciences Ross Lynn Fellowship (AB), support from the NIH Chemistry-Biology Interface Research Training program (T32 GM125620 to support NM), the Walther Cancer Foundation, the Ralph W. and Grace M. Showalter Research Trust (JH). In addition, we gratefully acknowledge Tripti Bera for technical support.

## Author Contributions

JH conceived the project. AB and JH were responsible for study design and supervision. AB, ET, NM, AM, BL, HM, MS, LG, CP, and NJ acquired the data. AB, ET, NM, MS, LG, SU, EP, NL, and JH were responsible for the data analysis, data interpretation, and reagents. AB, SU, NL and JH were responsible for the interpretation and visualization of the data. AB and JH drafted the manuscript. All authors were responsible for critical revision of the manuscript.

## Competing Interests

No competing interests declared.

## Data Availability Statement

The RNA-sequencing transcriptomic data is deposited in the Gene Expression Omnibus (GEO) under accession number GSE240260.

## Notes

No conflicts of interest declared

### Competing Interest Statement

The authors have declared no competing interest.

## REFERENCES

1 D’Angelo SP, Munhoz RR, Kuk D, Landa J, Hartley EW, Bonafede M et al. Outcomes of Systemic Therapy for Patients with Metastatic Angiosarcoma. Oncology 2015; 89: 205–214.

2 Fury MG, Antonescu CR, Van Zee KJ, Brennan MF, Maki RG. A 14-year retrospective review of angiosarcoma: clinical characteristics, prognostic factors, and treatment outcomes with surgery and chemotherapy. Cancer J Sudbury Mass 2005; 11: 241–247.

3 Painter CA, Jain E, Tomson BN, Dunphy M, Stoddard RE, Thomas BS et al. The Angiosarcoma Project: enabling genomic and clinical discoveries in a rare cancer through patient-partnered research. Nat Med 2020; 26: 181–187.

4 Chan JY, Lim JQ, Yeong J, Ravi V, Guan P, Boot A et al. Multiomic analysis and immunoprofiling reveal distinct subtypes of human angiosarcoma. J Clin Invest 2020; 130: 5833–5846.

5 Behjati S, Tarpey PS, Sheldon H, Martincorena I, Van Loo P, Gundem G et al. Recurrent PTPRB and PLCG1 mutations in angiosarcoma. Nat Genet 2014; 46: 376–379.

6 Espejo-Freire AP, Elliott A, Rosenberg A, Costa PA, Barreto-Coelho P, Jonczak E et al. Genomic Landscape of Angiosarcoma: A Targeted and Immunotherapy Biomarker Analysis. Cancers 2021; 13: 4816.

7 Wagner MJ, Lyons YA, Siedel JH, Dood R, Nagaraja AS, Haemmerle M et al. Combined VEGFR and MAPK pathway inhibition in angiosarcoma. Sci Rep 2021; 11: 9362.

8 Grilley-Olson JE, Allred JB, Schuetze S, Davis EJ, Wagner MJ, Poklepovic AS et al. A multicenter phase II study of cabozantinib + nivolumab for patients (pts) with advanced angiosarcoma (AS) previously treated with a taxane (Alliance A091902). J Clin Oncol 2023; 41: 11503–11503.

9 Saha J, Kim JH, Amaya CN, Witcher C, Khammanivong A, Korpela DM et al. Propranolol Sensitizes Vascular Sarcoma Cells to Doxorubicin by Altering Lysosomal Drug Sequestration and Drug Efflux. Front Oncol 2021; 10.https://www.frontiersin.org/articles/10.3389/fonc.2020.614288 (accessed 12 Sep2023).

10 Abraham JA, Hornicek FJ, Kaufman AM, Harmon DC, Springfield DS, Raskin KA et al. Treatment and Outcome of 82 Patients with Angiosarcoma. Ann Surg Oncol 2007; 14: 1953–1967.

11 Buehler D, Rice SR, Moody JS, Rush P, Hafez G-R, Attia S et al. Angiosarcoma outcomes and prognostic factors: a 25-year single institution experience. Am J Clin Oncol 2014; 37: 473–479.

12 Bartel DP. MicroRNAs: Target Recognition and Regulatory Functions. Cell 2009; 136: 215–233.

13 Adams BD, Kasinski AL, Slack FJ. Aberrant Regulation and Function of MicroRNAs in Cancer. Curr Biol 2014; 24: R762–R776.

14 Calin GA, Dumitru CD, Shimizu M, Bichi R, Zupo S, Noch E, et al. Frequent deletions and down-regulation of micro-RNA genes miR15 and miR16 at 13q14 in chronic lymphocytic leukemia. Proc Natl Acad Sci 2002; 99: 15524–15529.

15 Cimmino A, Calin GA, Fabbri M, Iorio MV, Ferracin M, Shimizu M et al. miR-15 and miR-16 induce apoptosis by targeting BCL2. Proc Natl Acad Sci 2005; 102: 13944– 13949.

16 Medina PP, Nolde M, Slack FJ. OncomiR addiction in an in vivo model of microRNA-21-induced pre-B-cell lymphoma. Nature 2010; 467: 86–90.

17 Chang T-C, Yu D, Lee Y-S, Wentzel EA, Arking DE, West KM et al. Widespread microRNA repression by Myc contributes to tumorigenesis. Nat Genet 2008; 40: 43– 50.

18 Hanna JA, Garcia MR, Go JC, Finkelstein D, Kodali K, Pagala V et al. PAX7 is a required target for microRNA-206-induced differentiation of fusion-negative rhabdomyosarcoma. Cell Death Dis 2016; 7: e2256.

19 Svoronos AA, Engelman DM, Slack FJ. OncomiR or Tumor Suppressor? The Duplicity of MicroRNAs in Cancer. Cancer Res 2016; 76: 3666–3670.

20 Hanna JA, Garcia MR, Lardennois A, Leavey PJ, Maglic D, Fagnan A et al. PAX3-FOXO1 drives miR-486-5p and represses miR-221 contributing to pathogenesis of alveolar rhabdomyosarcoma. Oncogene 2018; 37: 1991–2007.

21 Orellana EA, Tenneti S, Rangasamy L, Lyle LT, Low PS, Kasinski AL. FolamiRs: Ligand-targeted, vehicle-free delivery of microRNAs for the treatment of cancer. Sci Transl Med 2017; 9. doi:10.1126/scitranslmed.aam9327.

22 Rupaimoole R, Slack FJ. MicroRNA therapeutics: towards a new era for the management of cancer and other diseases. Nat Rev Drug Discov 2017; 16: 203– 222.

23 La Ferlita A, Sp N, Goryunova M, Nigita G, Pollock RE, Croce CM et al. Small Non-Coding RNAs in Soft-Tissue Sarcomas: State of the Art and Future Directions. Mol Cancer Res 2023; 21: 511–524.

24 Nakashima S, Jinnin M, Kanemaru H, Kajihara I, Igata T, Okamoto S et al. The role of miR-210, E2F3 and ephrin A3 in angiosarcoma cell proliferation. Eur J Dermatol 2017; 27: 464–471.

25 Heishima K, Mori T, Sakai H, Sugito N, Murakami M, Yamada N et al. MicroRNA-214 Promotes Apoptosis in Canine Hemangiosarcoma by Targeting the COP1-p53 Axis. PLoS ONE 2015; 10. doi:10.1371/journal.pone.0137361.

26 Wang X, Song Y. MicroRNA-340 inhibits the growth and invasion of angiosarcoma cells by targeting SIRT7. Biomed Pharmacother Biomedecine Pharmacother 2018; 103: 1061–1068.

27 Chen Y, Kuang D, Zhao X, Chen D, Wang X, Yang Q et al. miR-497-5p inhibits cell proliferation and invasion by targeting KCa3.1 in angiosarcoma. Oncotarget 2016; 7: 58148–58161.

28 Loh JW, Lee JY, Lim AH, Guan P, Lim BY, Kannan B, et al. Spatial transcriptomics reveal topological immune landscapes of Asian head and neck angiosarcoma. Commun Biol 2023; 6: 1–9.

29 Hanna JA, Drummond CJ, Garcia MR, Go JC, Finkelstein D, Rehg JE et al. Biallelic Dicer1 Loss Mediated by aP2-Cre Drives Angiosarcoma. Cancer Res 2017; 77: 6109–6118.

30 Hanna JA, Langdon CG, Garcia MR, Benton A, Lanman NA, Finkelstein D et al. Genetic context of oncogenic drivers dictates vascular sarcoma development in aP2-Cre mice. J Pathol 2022. doi:10.1002/path.5873.

31 Nagy Á, Munkácsy G, Győrffy B. Pancancer survival analysis of cancer hallmark genes. Sci Rep 2021; 11: 6047.

32 Hadj-Hamou N-S, Laé M, Almeida A, Grange P de la, Kirova Y, Sastre-Garau X et al. A transcriptome signature of endothelial lymphatic cells coexists with the chronic oxidative stress signature in radiation-induced post-radiotherapy breast angiosarcomas. Carcinogenesis 2012; 33: 1399–1405.

33 Lewis BP, Burge CB, Bartel DP. Conserved Seed Pairing, Often Flanked by Adenosines, Indicates that Thousands of Human Genes are MicroRNA Targets. Cell 2005; 120: 15–20.

34 Wu R, Tang S, Wang M, Xu X, Yao C, Wang S. MicroRNA-497 Induces Apoptosis and Suppresses Proliferation via the Bcl-2/Bax-Caspase9-Caspase3 Pathway and Cyclin D2 Protein in HUVECs. PloS One 2016; 11: e0167052–e0167052.

35 Gleissner CM-L, Pyka CL, Heydenreuter W, Gronauer TF, Atzberger C, Korotkov VS et al. Neocarzilin A Is a Potent Inhibitor of Cancer Cell Motility Targeting VAT-1 Controlled Pathways. ACS Cent Sci 2019; 5: 1170–1178.

36 Heng W, Wei F, Li W, Li Q, Xiong D-L, Ma Y-Y et al. Expression of VAT1 in hepatocellular carcinoma and its clinical significance. Neoplasma 2021; 68: 416– 422.

37 Shan X, Wang K, Tong X, Wang Z, Wu F, Liu X et al. High expression of VAT1 is a prognostic biomarker and predicts malignancy in glioblastoma. Oncol Rep 2019; 42: 1422–1430.

38 Mertsch S, Becker M, Lichota A, Paulus W, Senner V. Vesicle amine transport protein-1 (VAT-1) is upregulated in glioblastomas and promotes migration. Neuropathol Appl Neurobiol 2009; 35: 342–352.

39 Junker M, Rapoport TA. Involvement of VAT-1 in Phosphatidylserine Transfer from the Endoplasmic Reticulum to Mitochondria. Traffic 2015; 16: 1306–1317.

40 Watanabe Y, Tamura Y, Kakuta C, Watanabe S, Endo T. Structural basis for interorganelle phospholipid transport mediated by VAT-1. J Biol Chem 2020; 295: 3257–3268.

41 Kim S-Y, Mori T, Chek MF, Furuya S, Matsumoto K, Yajima T et al. Structural insights into vesicle amine transport-1 (VAT-1) as a member of the NADPH-dependent quinone oxidoreductase family. Sci Rep 2021; 11: 2120.

42 Yang P, Wang K, Zhang C, Wang Z, Liu Q, Wang J et al. Novel roles of VAT1 expression in the immunosuppressive action of diffuse gliomas. Cancer Immunol Immunother 2021; 70: 2589–2600.

43 Drummond CJ, Hanna JA, Garcia MR, Devine DJ, Heyrana AJ, Finkelstein D et al. Hedgehog Pathway Drives Fusion-Negative Rhabdomyosarcoma Initiated From Non-myogenic Endothelial Progenitors. Cancer Cell 2018; 33: 108–124.e5.

44 Forterre A, Komuro H, Aminova S, Harada M. A Comprehensive Review of Cancer MicroRNA Therapeutic Delivery Strategies. Cancers 2020; 12: 1852.

45 Orellana E, Kasinski A. MicroRNAs in Cancer: A Historical Perspective on the Path from Discovery to Therapy. Cancers 2015; 7: 1388–1405.

46 Boichard A, Wagner MJ, Kurzrock R. Angiosarcoma heterogeneity and potential therapeutic vulnerability to immune checkpoint blockade: insights from genomic sequencing. Genome Med 2020; 12: 61.

47 Chen S, Fu Z, Wen S, Yang X, Yu C, Zhou W et al. Expression and Diagnostic Value of miR-497 and miR-1246 in Hepatocellular Carcinoma. Front Genet 2021; 12.https://www.frontiersin.org/articles/10.3389/fgene.2021.666306 (accessed 22 Jul2023).

48 Li D, Zhao Y, Liu C, Chen X, Qi Y, Jiang Y et al. Analysis of MiR-195 and MiR-497 Expression, Regulation and Role in Breast Cancer. Clin Cancer Res 2011; 17: 1722–1730.

49 Feng L, Cheng K, Zang R, Wang Q, Wang J. miR-497-5p inhibits gastric cancer cell proliferation and growth through targeting PDK3. Biosci Rep 2019; 39: BSR20190654.

50 Davis AJ, Tsinkevich M, Rodencal J, Abbas HA, Su X, Gi Y-J et al. TAp63-Regulated miRNAs Suppress Cutaneous Squamous Cell Carcinoma through Inhibition of a Network of Cell-Cycle Genes. Cancer Res 2020; 80: 2484–2497.

51 Lobov I, Mikhailova N. The Role of Dll4/Notch Signaling in Normal and Pathological Ocular Angiogenesis: Dll4 Controls Blood Vessel Sprouting and Vessel Remodeling in Normal and Pathological Conditions. J Ophthalmol 2018; 2018: 3565292.

52 Hellström M, Phng L-K, Hofmann JJ, Wallgard E, Coultas L, Lindblom P et al. Dll4 signalling through Notch1 regulates formation of tip cells during angiogenesis. Nature 2007; 445: 776–780.

53 Nozoe S, Ishii N, Kusano G, Kikuchi K, Ohta T. Neocarzilins A and B, Novel Polyenones from Streptomyces Carzinostaticus. Tetrahedron Lett 1992; 33: 7547–7550.

54 Italiano A, Chen C-L, Thomas R, Breen M, Bonnet F, Sevenet N et al. Alterations of the p53 and PIK3CA/AKT/mTOR pathways in angiosarcomas: a pattern distinct from other sarcomas with complex genomics. Cancer 2012; 118: 5878–5887.

55 Gorden BH, Saha J, Khammanivong A, Schwartz GK, Dickerson EB. Lysosomal drug sequestration as a mechanism of drug resistance in vascular sarcoma cells marked by high CSF-1R expression. Vasc Cell 2014; 6: 20.

56 Chen S. Ultrafast one-pass FASTQ data preprocessing, quality control, and deduplication using fastp. iMeta 2023; 2: e107.

57 Dobin A, Davis CA, Schlesinger F, Drenkow J, Zaleski C, Jha S et al. STAR: ultrafast universal RNA-seq aligner. Bioinformatics 2013; 29: 15.

58 Liao Y, Smyth GK, Shi W. featureCounts: an efficient general purpose program for assigning sequence reads to genomic features. Bioinformatics 2014; 30: 923–930.

59 Robinson MD, McCarthy DJ, Smyth GK. edgeR: a Bioconductor package for differential expression analysis of digital gene expression data. Bioinformatics 2010; 26: 139–140.

60 Dennis G, Sherman BT, Hosack DA, Yang J, Gao W, Lane HC et al. DAVID: Database for Annotation, Visualization, and Integrated Discovery. Genome Biol 2003; 4: R60.

61 Fellmann C, Hoffmann T, Sridhar V, Hopfgartner B, Muhar M, Roth M et al. An Optimized microRNA Backbone for Effective Single-Copy RNAi. Cell Rep 2013; 5: 1704–1713.

62 Carson FL, Cappellano CH. Histotechnology. A self instructional text. 4th edition. ASCP: Chicago, 2015.

