## Supplementary Material for "Target gene regulatory network of miR-497 in angiosarcoma"

### Supplementary Information Inventory

**Supplementary Methods**
**Supplementary Figure Legends**
**Supplementary Figures**

Figure S1. Validation of microRNA mimic transfection in cell line panel.
Figure S2. Kaplan-Meier survival plot of miR-497 expression in sarcomas.
Figure S3. MicroRNA target gene enrichment in miR-497 mimic transfected cells.
Figure S4. Validation of miR-497 target genes.
Figure S5. Knockdown of miR-497 target gene, VAT1, inhibits cell migration.

**Supplementary Tables**

Table S1. miR-497 family predicted target genes significantly downregulated with miR-497 mimic.
Table S2. Gene Ontology analysis of miR-497 upregulated genes in ADC106 cells.
Table S3. Gene Ontology analysis of miR-497 downregulated genes in ADC106 cells.
Table S4. Gene ontology analysis of common miR-497 mimic downregulated genes and upregulated genes in AD$^{cKO}$ tumors.
Table S5. STR Validation of cell lines.
Table S6. SYBR and miRCURY primers used for qRT-PCR.
Table S7. Primers for pre-miR-497 and 3` UTR cloning of miR-497 target genes.
Table S8. Antibodies used for immunoblots.

### Supplementary Methods

#### Molecular Cloning

All cloning primers are detailed in Supplementary Table S7. The 749 base pair genomic fragment of human pre-miR-497-195 was PCR amplified from pMSCV-PIG-miR-497/miR-195 (Addgene #64236, a gift from Joshua Mendell) and subcloned into the pCMV6 expression plasmid using the EcoRI and SalI sites to generate pCMV6-miR-497/195. The sequence for mClover was purchased as a gene block from IDT and cloned into pSIN-puro using the SpeI restriction site, generating pSIN-mClover-IRES-Puro. The 292 base pair fragment containing just the pre-miR-497 was PCR amplified from pCMV6-miR-497/195 and cloned into pSIN-mClover using the EcoRI site, generating pSIN-mClover-pre-miR-497-IRES-Puro. The miR-497 sensor with the perfect reverse complement sequence of miR-497 was constructed by annealing oligos and ligating them into XhoI and NotI digested psiCHECK2 miR-497 in psiCHECK2 using (C8021; Promega). The *Ccnd2*, *Dll4*, and *Cdk6* 3` UTRs were PCR amplified from ADC106 or AS5 genomic DNA for mouse and human 3` UTRs, respectively. The 3`UTR of Vat1 was purchased as a gene block from IDT (encoding the sequence from the stop codon to nucleotide 692 of 3` UTR). 3` UTRs were cloned into the psiCHECK2 reporter plasmid using Xho and NotI restriction sites primarily using Gibson Assembly (E2621S, New England Biolabs, Ipswich, MA). The miR-497 recognition site in the Vat1 3` UTR was mutated with the Quick Change II Site Directed Mutagenesis (200524; Agilent Technologies, Santa Clara, CA, USA). The mutated 3` UTRs of *Dll4* and *Cdk6* were purchased as gene blocks from IDT (miR-497 sites changed from TGCT**GCT** changed to TGCT**TAC**) and cloned into psiCHECK2 using Xho and NotI restriction sites.

#### Chemical Synthesis of Neocarzilin A

All reagents and solvents were purchased reagent grade or higher from commercial vendors (Sigma-Aldrich, Thermo Fisher Scientific Inc., Oakwood Chemical, Alfa Aesar, Acros Chemicals) and were used without further purification. All reactions were conducted using flame dried glassware whereas reactions containing oxygen or water sensitive reagents were carried under nitrogen atmosphere.

For reaction monitoring, analytical thin-layer chromatography (TLC) was carried out on Merck silica-gel 60 F254 plates, using short wave UV light (λ=254 nm) or KMnO4-stain (1.50 g KMnO4, 10.0 g K2CO3, 1.25 mL NaOHaq (10 wt-%), 200 mL ddH2O) to visualize reaction components.

1H-NMR experiments were recorded on Avance-III (AV-HD400 or DRX-HD500) NMR systems (Bruker Co.) at room temperature with CDCl3 as solvents and referenced to the residual proton signal of the corresponding deuterated solvent (CDCl3: δ = 7.26 ppm). Chemical shifts are reported in parts per million (ppm). Coupling constants (J) are reported in hertz (Hz) and for the assignment of multiplicity to the signals the following abbreviations were used: s = singlet, d = doublet, t = triplet, q = quartet, m = multiplet or unresolved.

**Ethyl (E)-4-(diethoxyphosphoryl)-but-2-enoate**

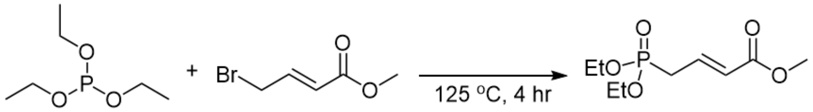


Triethyl phosphite (5.1 g, 30.7 mmol, 1.1 equiv.) was added to methyl 4-bromo-crotonate (5.0 g, 27.9 mmol, 1.0 equiv.) and heated to a reflux at 130 °C for 4 hours. The crude product was purified via vacuum distillation (115 °C, 0.04 mbar). 6.2 g (26.2 mmol, 94%) of desired product **1** was obtained as a yellow oil.

**$^1$H NMR** (400 MHz, $CDCl_3$) δ (ppm) = 6.94 – 6.82 (m, 1H), 5.96 (ddt, J = 15.6, 5.0, 1.7 Hz, 1H), 4.18 – 4.06 (m, 4H), 3.74 (s, 3H), 2.74 (ddd, J = 22.9, 7.9, 1.4 Hz, 2H), 1.32 (d, J = 7.1 Hz, 6H).

**Methyl (S,2E,4E)-6-methylocta-2,4-dienoate**

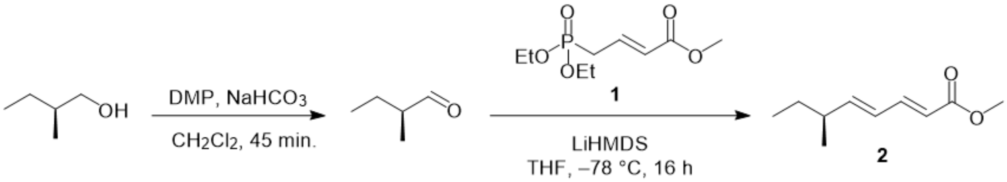


Dess-Martin Periodinane reagent (6 g, 14.18 mmol, 1.25 equiv.) and $NaHCO_3$ (0.25 g, 2.80 mmol, 0.25 equiv.) were dissolved in 18mL of $CH_2Cl_2$ in a flame dried flask and cooled to 0 °C in an ice bath. S-2-methyl-butan-1-ol (1 g, 11.34 mmol, 1.0 equiv.) was added slowly and stirred at 0 °C for 45 minutes. Diethyl ether (20mL) was added and the reaction was quenched by slow addition of NaOH (3.75 M, 20mL). The resulting suspension was allowed to stir a further 15 minutes until a clear solution was reached, upon which the organics were separated and the aqueous was washed with diethyl ether (10mL). The organics were combined and dried with $Na_2SO_4$, filtered, and the organics were removed under a stream of $N_2$. In parallel, **1** (3.75 g, 15.90 mmol, 1.4 equiv.) was dissolved in anhydrous THF (12mL) and cooled to -78 °C. LiHMDS (1M in THF, 17.01 mL, 17.01 mmol, 1.5 equiv.) was added slowly and the reaction stirred for 1 hr at -78 °C followed by addition of the aldehyde. The resulting mixture was stirred for 1 hr, upon which it was allowed to warm to -40 °C and stir a further 4 hours. The reaction was then diluted with diethyl ether (50mL) and allowed to warm to 0 °C and quenched by addition of sat. aq. $NH_4Cl$ (50mL). The organics were washed further with sat. aq. $NH_4Cl$ (3 x 25 mL), dried with $Na_2SO_4$, filtered, and concentrated. The crude product was purified via flash chromatography to afford **2** (1.33 g, 70% over two steps) as a yellow oil.

**Rf =** 0.25 (3% EtOAc in Hexanes) [UV, $KMnO_4$]
**$^1$H NMR** (400 MHz, $CDCl_3$) δ (ppm) = 7.26 (dd, J = 15.7, 10.6 Hz, 1H), 6.19 – 5.96 (m, 2H), 5.80 (d, J = 15.4 Hz, 1H), 3.73 (s, 3H), 2.16 (hept, 1H), 1.37 (d, J = 7.5 Hz, 2H), 1.03 (d, J = 6.7 Hz, 3H), 0.86 (t, J = 7.1 Hz, 3H).

**(S,2E,4E)-6-methylocta-2,4-dien-1-ol**

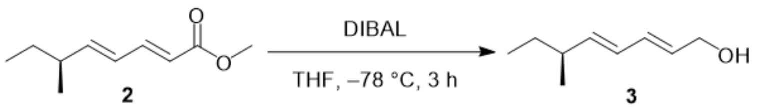


Methyl ester **2** (0.50 g, 2.97 mmol, 1 equiv.) was dissolved in anhydrous THF (7mL) and cooled to -78 °C. DIBAL-H (1.1M in cyclohexane, 6.76mL, 2.5 equiv.) was added slowly and allowed to react for 3 hours. The reaction was allowed to warm to 0 °C and diluted with diethyl ether (20 mL). The reaction was quenched by the addition of sat. aq. potassium sodium tartrate (20mL) and was stirred a further 15 minutes. The aqueous was washed with diethyl ether (20 mL) and the organics were dried with $Na_2SO_4$, filtered, and concentrated. The resulting colorless oil **3** (0.41 g, 97%) was sufficiently pure to use without purification.

**$^1$H NMR** (400 MHz, $CDCl_3$) δ (ppm) = 6.28 – 6.17 (m, 1H), 6.01 (dd, J = 15.2, 10.4 Hz, 1H), 5.74 (dt, J = 15.2, 6.1 Hz, 1H), 5.59 (dd, J = 15.2, 7.8 Hz, 1H), 4.20 – 4.12 (m, 2H), 2.07 (hept, J = 6.9 Hz, 1H), 1.38 – 1.29 (m, 2H), 0.99 (d, J = 6.7 Hz, 3H), 0.85 (t, J = 7.4 Hz, 3H).

**(S,3E,5E,7E)-9-methylundeca-3,5,7-trien-2-one**

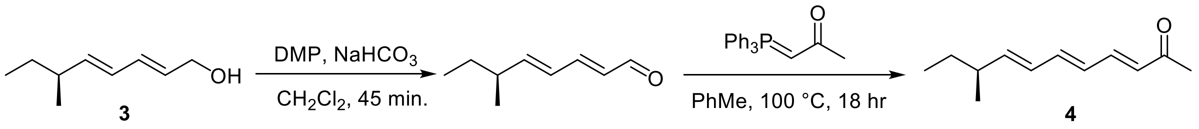


Dess-Martin Periodinane reagent (1.14 g, 2.68 mmol, 1.25 equiv.) and $NaHCO_3$ (45 mg, 0.535 mmol, 0.25 equiv.) were dissolved in anhydrous $CH_2Cl_2$ (10mL) in a flame dried flask. The mixture was cooled to 0 °C and alcohol **3** (0.30 g, 2.14 mmol, 1 equiv.) was added. The reaction was allowed to stir for 45 minutes and was diluted with diethyl ether (20mL) and quenched with addition of NaOH (3.75 M, 20 mL) and stirred for 10 minutes. The organics were washed with NaOH (1 M, 20 mL), dried with $Na_2SO_4$, filtered, and concentrated to a colorless oil. The aldehyde was dissolved in anhydrous toluene (15mL) and added to 1-(Triphenylphosphoranylidene)-2-propanone (1.36 g, 4.28 mmol, 2 equiv.) in a flame-dried flask. The reaction was heated near reflux and stirred for 18 hours at 100 °C. The reaction is then slowly cooled to 0 °C and cold hexanes are added to precipitate the triphenyl phosphine oxide byproduct. The TPPO is filtered off, the solvent evaporated, and the precipitation is repeated twice, or until sufficient TPPO is removed. The crude product is purified via flash chromatography to afford ketone **4** (0.20 g, 1.14 mmol, 53% over two steps) as a pale-yellow oil.

**Rf** = 0.32 (10% EtOAc in Hexanes) [UV, $KMnO_4$]

$^{1}$**H NMR** (400 MHz, $CDCl_3$) δ (ppm) = 7.14 (dd, J = 15.6, 11.1 Hz, 1H), 6.58 (dd, J = 14.9, 10.7 Hz, 1H), 6.24 (dd, J = 14.9, 11.1 Hz, 1H), 6.17 – 6.07 (m, 2H), 5.84 (dd, J = 15.2, 7.8 Hz, 1H), 2.27 (s, 3H), 2.15 (p, J = 7.0 Hz, 1H), 1.45 – 1.30 (m, J = 7.3, 6.7 Hz, 2H), 1.02 (d, J = 6.7 Hz, 3H), 0.87 (t, J = 7.4 Hz, 3H).

**(S,3Z,5E,7E,9E)-1,1,1-trichloro-4-hydroxy-11-methyltrideca-3,5,7,9-tetraen-2-one**

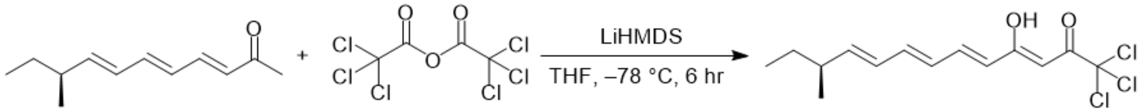


Ketone **4** (0.10 g, 0.56 mmol, 1 equiv.) was dissolved in anhydrous THF (5mL) in a flame dried flask and cooled to -78 °C. LiHMDS (1M in THF, 0.62 mL, 0.62 mmol, 1.1 equiv.) was added slowly and the reaction was allowed to stir at -78 °C for 1 hour. Trichloroacetic anhydride (0.21 mL, 1.12 mmol, 2 equiv.) was then added slowly and the reaction stirred for 5 hours at -78 °C. The reaction was diluted with diethyl ether (20 mL), allowed to warm to 0 °C, and was quenched by the addition of sat. aq. $NH_4Cl$ (20 mL), and allowed to warm to rt. The organics were washed with sat. aq. $NaHCO_3$ (3 x 20 mL), sat. aq. $NH_4Cl$ (3 x 20 mL), and brine (2 x 20 mL). The organics were dried with $Na_2SO_4$, filtered, and concentrated to a bright yellow oil. The crude product was purified via flash chromatography to afford Neocarzilin A (135mg, 0.42 mmol, 75%) as an orange oil.

**Rf =** 0.38 (5% EtOAc in Hexanes) [UV, $KMnO_4$]
$^{1}$**H NMR** (400 MHz, $CDCl_3$) δ (ppm) = 7.37 (ddd, J = 14.9, 11.5, 0.5 Hz, 1H), 6.62 (dd, J = 14.8, 10.7 Hz, 1H), 6.27 (dd, J = 14.8, 11.3 Hz, 1H), 6.21 – 6.08 (m, 2H), 6.00 (d, J = 15.0 Hz, 1H), 5.89 (dd, J = 15.2, 7.8 Hz, 1H), 2.17 (p, J = 6.9 Hz, 1H), 1.45 – 1.31 (m, 2H), 1.03 (d, J = 6.7 Hz, 3H), 0.87 (t, J = 7.4 Hz, 3H).
$^{13}$**C NMR** ($CDCl_3$) δ (ppm) = 186.0, 177.9, 147.6, 142.9, 142.7, 128.3, 128.3, 123.2, 95.0, 93.0, 38.8, 29.4, 19.6, 11.6
**HRMS ESI** calcd. for $C_{14}H_{17}Cl_3O_2$ [M+H]: 323.0294, found: 323.0368.
$[\alpha]^{22}_{D}$ = 49.6 (c = 1.0, $CHCl_3$)

**Supplementary Figure Legends**

**Figure S1. Validation of microRNA mimic transfection in cell line panel. (A)** Relative expression of miR-23-3p, **(B)** miR-210-3p, **(C)** miR-214-3p, and **(D)** miR-340-5p by qRT-PCR in cells as in mouse and human cells, 72 hours post-transfection with indicated miRNA mimics relative to negative control (NC) mimic. *p<0.05.

**Figure S2. Kaplan-Meier survival plot of miR-497 expression in sarcomas. (A)** Relative cell viability of pSIN-Clover-empty control and pSIN-Clover-pre-miR-497 ADC106 cells. **(B)** Kaplan-Meier survival plot of sarcoma patients (n=259) from the SARC TCGA dataset using Kaplan-Meier Plotter (Lanczky et al. 2021).

**Figure S3. MicroRNA target gene enrichment in miR-497 mimic transfected cells. (A)** Heatmap of genes significantly up or downregulated upon miR-497 mimic transfection **(B)** Enrichment of downregulated microRNA target genes in ADC106 cells transfected with miR-497 mimic. Genes downregulated with p<0.05 and log FC > 1 entered into Enrichr (846 genes). **(C)** Gene ontology analysis with significantly enriched biological pathways in genes upregulated in cells transfected with miR-497 mimic from Fig 4A with p < 0.05 and log FC > 1 (559 genes). **(D)** Gene ontology analysis with significantly enriched pathways in genes downregulated in cells transfected with miR-497 mimic from Fig 4A with P < 0.05 and log FC < -1 (846 genes).

**Figure S4. Validation of miR-497 target genes. (A)** Mapped miR-497 binding sites in the 3` UTRs of candidate target genes, *Ccnd2*, **(B)** *Cdk6*, and **(C)** *Vat1*. **(D)** Luciferase activity in 293T cells co-transfected with pre-miR-195/497 or control vector, and empty psiCHECK2 reporter, miR-497 sensor positive control, mouse wild-type human CDK6 3′ UTR, and 3` UTR with mutated miR-497 site. Renilla/Firefly luciferase ratio normalized to empty reporter (no miR-497), *p<0.05.

**Figure S5: Knockdown of miR-497 target gene, VAT1, inhibits cell migration. (A)** qRT-PCR and **(B)** immunoblot validation of *Vat1* expression in ADC106 and **(C)** SVR cells after 3 and 5 days with indicated doxycycline concentration treatments. **(C)** Transwell migration assays performed in ADC106 and SVR cells with control and Vat1 *shRNA* knockdown, **(D)** Quantification of the number of migrated cells as from (C). *p<0.05, **p<0.01, ***p<0.001, ****p<0.0001.

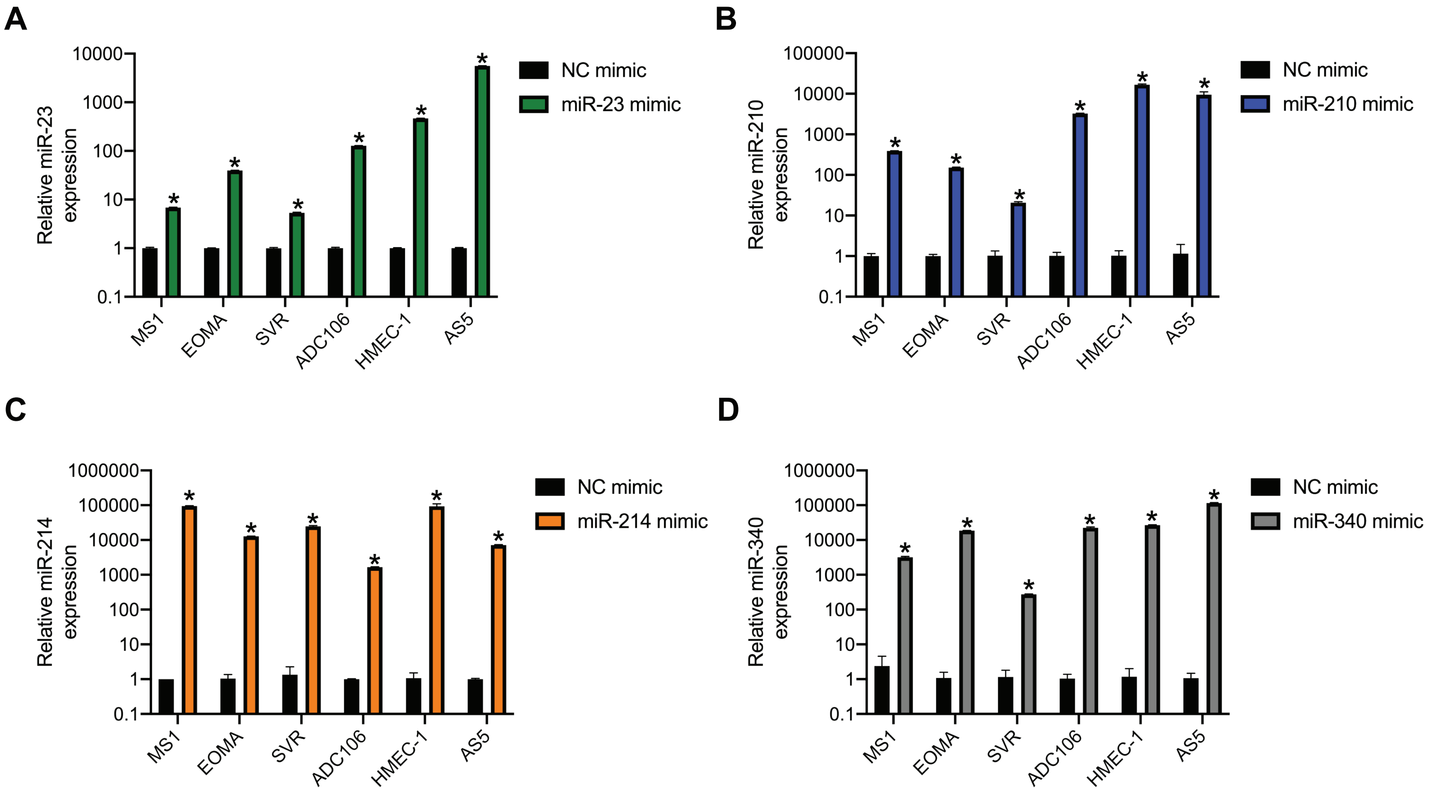


**Supplementary Figure 1.**

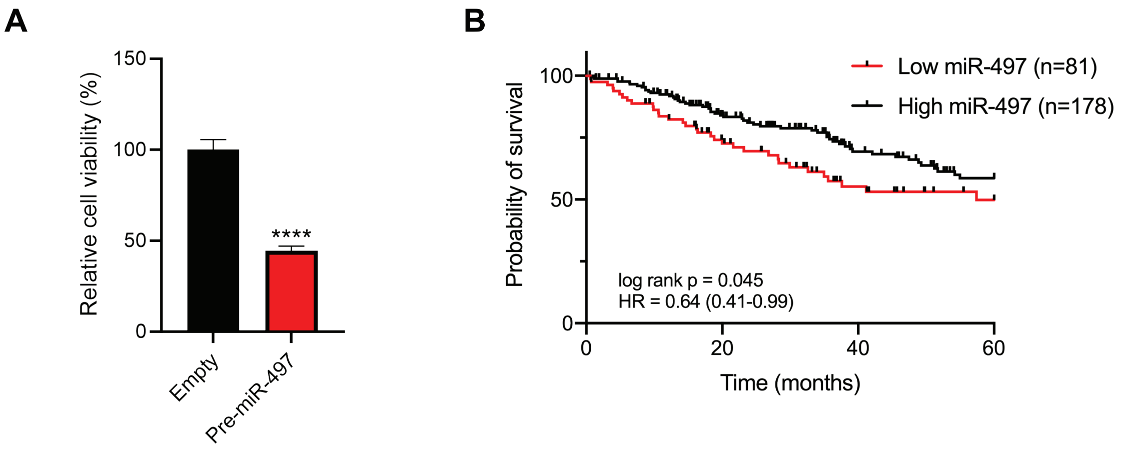


Supplementary Figure 2

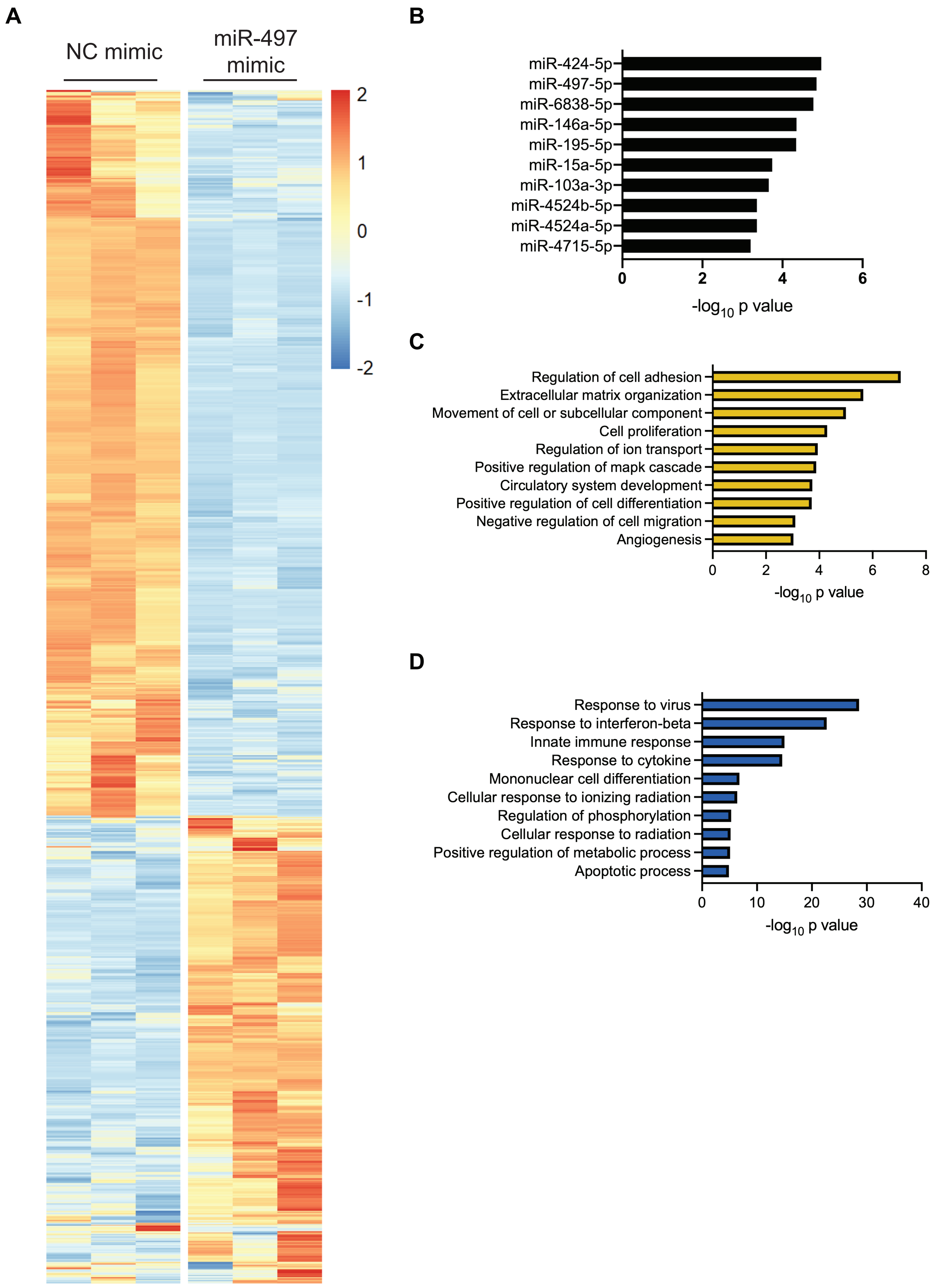


**Supplementary Figure 3.**

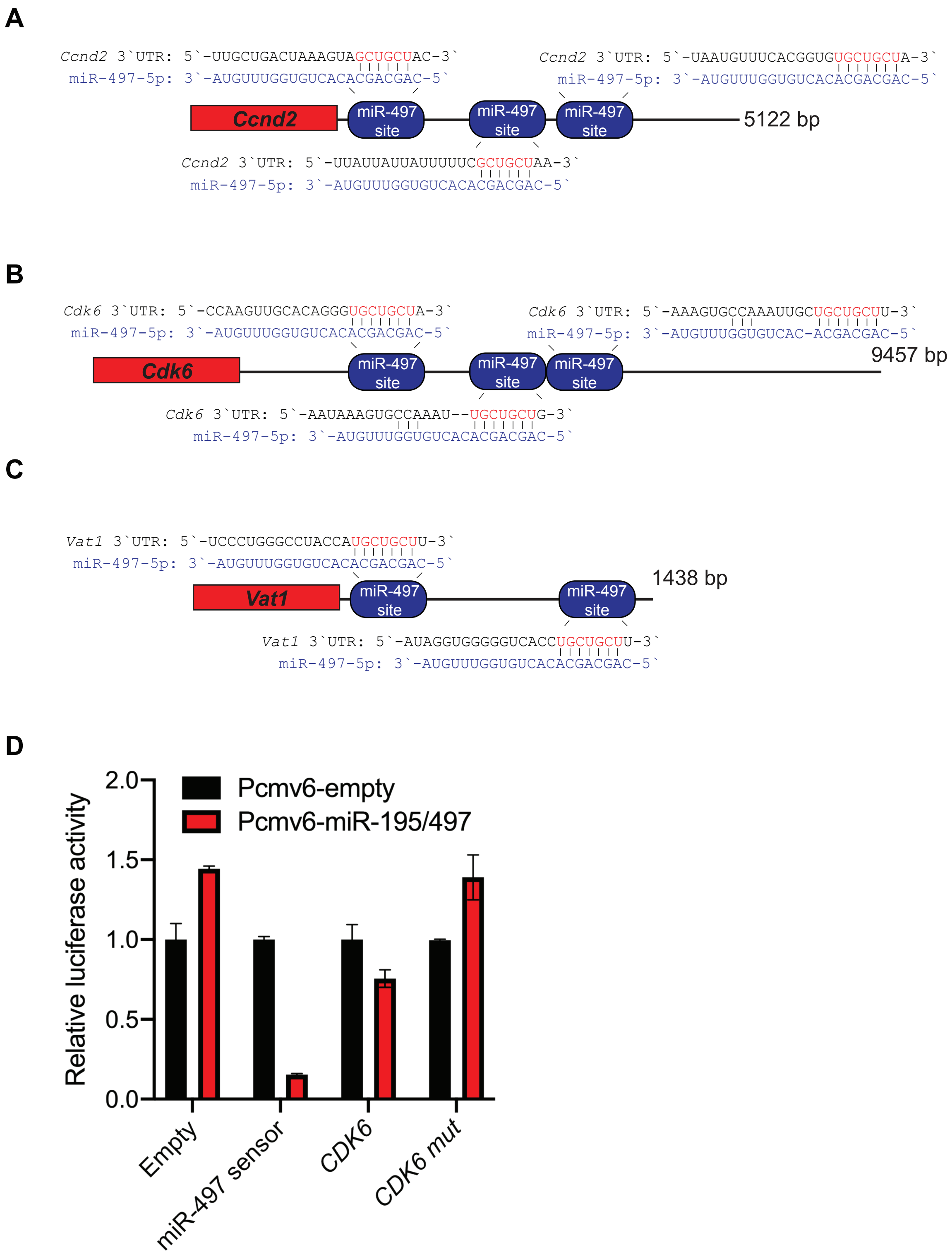


**Supplementary Figure 4.**

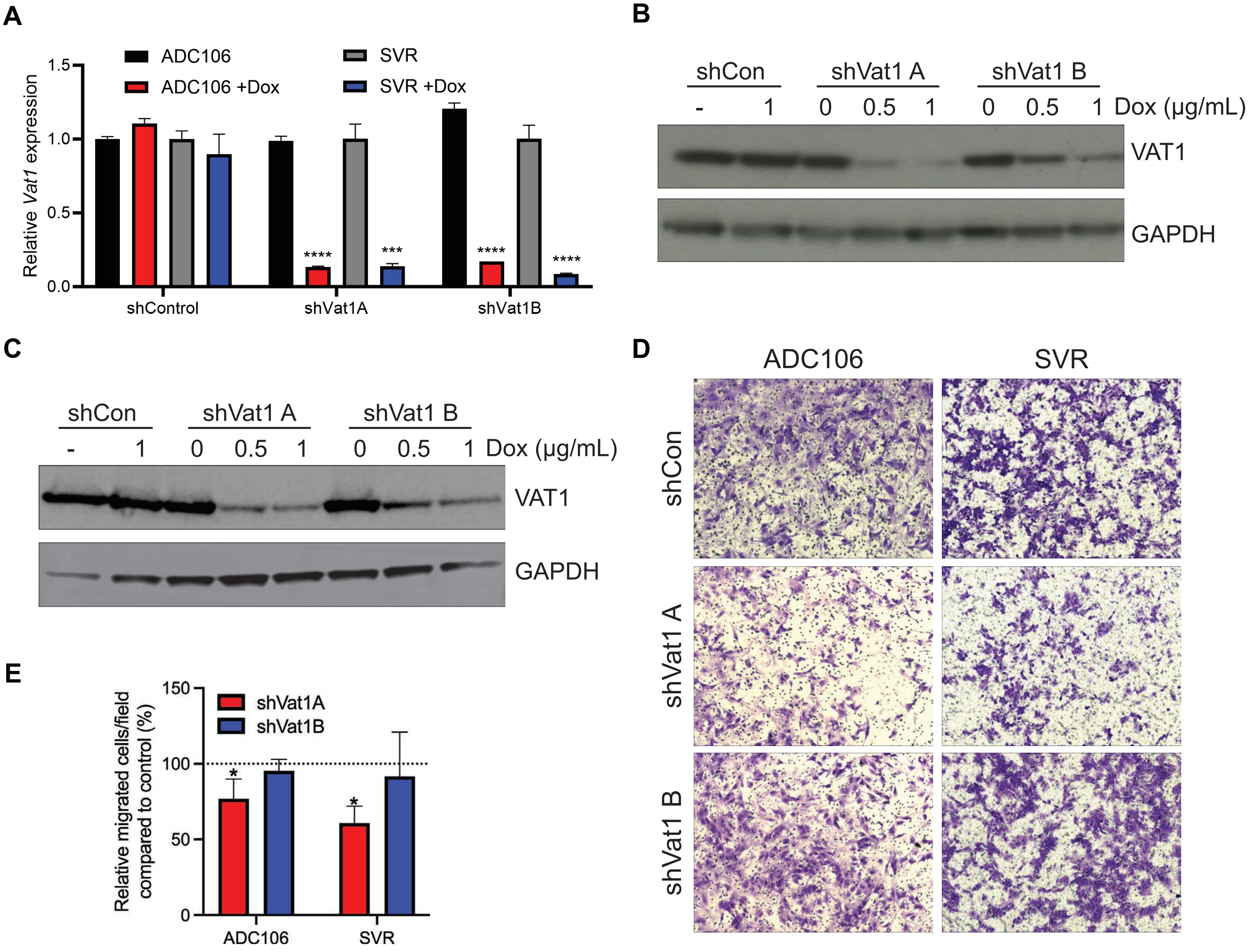


**Supplementary Figure 5**
